# Perception and Memory Retrieval States are Reflected in Distributed Patterns of Background Functional Connectivity

**DOI:** 10.1101/2022.09.14.507854

**Authors:** Y. Peeta Li, Yida Wang, Nicholas B. Turk-Browne, Brice A. Kuhl, J. Benjamin Hutchinson

## Abstract

The same visual input can serve as the target of perception or as a trigger for memory retrieval depending on whether cognitive processing is externally oriented (perception) or internally oriented (memory retrieval). While numerous human neuroimaging studies have characterized how visual stimuli are differentially processed during perception versus memory retrieval, perception and memory retrieval may also be associated with distinct neural states that are independent of stimulus-evoked neural activity. Here, we combined human fMRI with full correlation matrix analysis (FCMA) to reveal potential differences in “background” functional connectivity across perception and memory retrieval states. We found that perception and retrieval states could be discriminated with high accuracy based on patterns of connectivity across (1) the control network, (2) the default mode network (DMN), and (3) retrosplenial cortex (RSC). In particular, clusters in the control network increased connectivity with each other during the perception state, whereas clusters in the DMN were more strongly coupled during the retrieval state. Interestingly, RSC switched its coupling between networks as the cognitive state shifted from retrieval to perception. Finally, we show that background connectivity 1) was fully independent from stimulus-related variance in the signal and, further, 2) captured distinct aspects of cognitive states compared to traditional classification of stimulus-evoked responses. Together, our results reveal that perception and memory retrieval are associated with sustained cognitive states that manifest as distinct patterns of connectivity among large-scale brain networks.

## INTRODUCTION

When a familiar visual stimulus (e.g., a colleague) is encountered, it might serve as the target of perceptual scrutiny (e.g., their current facial expression) or the trigger for episodic memory retrieval (e.g., something they said in the past). Numerous cognitive theories and neuroimaging studies have contrasted the cognitive processes and brain regions engaged during perception versus retrieval (Bosch et al., 2014; Chun & Johnson, 2011; Kosslyn et al., 1995; McClelland et al., 1995; O’Reilly & McClelland, 1994; Polyn et al., 2005; Wheeler et al., 2000). The majority of this work has focused on investigating the operations performed upon the *stimulus*, such as how the stimulus is encoded into long-term memory or how the stimulus is used to drive memory recollection (Favila et al., 2020; Fernandez et al., 2022; Kim, 2013). However, evidence also suggests that perception and memory retrieval are characterized by distinct ‘states’ that are sustained over time and independent from external stimuli (Duncan et al., 2012; Duncan & Shohamy, 2016; Ezzyat et al., 2017; Guderian et al., 2009; Hasselmo et al., 1996). Characterizing how perception and retrieval states are implemented in the brain represents an important challenge, but one that is complicated by methodological and analytical factors.

One potentially powerful way to characterize distinct cognitive states is by measuring patterns of functional connectivity within the brain (Cohen & D’Esposito, 2016; Fritch et al., 2021; Shirer et al., 2012; Song & Rosenberg, 2021). Functional connectivity is typically computed over extended windows of time and is therefore well-suited to measure states that putatively persist across, or in the absence of, external stimuli. However, a critical issue in applying functional connectivity during cognitive tasks is that any observed correlations in neural activity may be largely or entirely driven by stimulus-evoked neural responses. That is, if two brain regions consistently respond to external stimuli, this will induce apparent “connectivity” between these regions. Several fMRI studies have addressed this concern by using “background” connectivity—an approach in which stimulus-evoked responses are explicitly modeled and removed, with connectivity then computed using the residual (background) activity (Al-Aidroos et al., 2012; Cole et al., 2019; Duncan et al., 2014; Norman-Haignere et al., 2012; Turk-Browne, 2013). Conceptually, this approach can isolate interactions between brain regions that reflect sustained, endogenous processing as opposed to transient, exogenous responses to stimuli (Summerfield et al., 2006; Turk-Browne, 2013).

To date, only a limited number of fMRI studies have used background connectivity to test for differences between perception and memory retrieval states (Cooper & Ritchey, 2019; Duncan et al., 2014). Moreover, these studies have only considered potential interactions (connectivity) between a relatively limited number of brain regions. For example, Cooper and Ritchey (2019) examined background FC patterns exclusively between *a priori*, memory-related brain regions in the posterior-medial (PM) and anterior-temporal (AT) networks. They found that regions in these networks exhibited stronger background connectivity during retrieval compared to perception, providing important evidence that perception versus retrieval states can be linked to stimulus-independent processes. While this type of targeted, seed-based analysis is highly valuable for testing specific hypotheses about regions of a priori interest, it is inherently blind to interactions involving non-seed regions (Turk-Browne, 2013). In other words, discoveries are systematically biased to come from the regions that are tested (Wang et al., 2015). Thus, there is value to unbiased approaches that allow for background FC to be more comprehensively measured—ideally, across the whole brain. That said, while conducting whole-brain functional connectivity may be theoretically appealing, it can be computationally intractable, particularly if applied without down-sampling data. For example, brain volumes consisting of 50,000 voxels would yield 1.25B voxel pairs (‘connections’) to analyze. For this reason, previous attempts to measure whole-brain functional connectivity have tended to substantially reduce the dimensionality of data by grouping voxels into regions or parcels (e.g., Pantazatos et al., 2012; Shirer et al., 2012; Watanabe et al., 2012).

Here, we sought to identify patterns of background connectivity associated with perception versus retrieval states using full correlation matrix analysis (FCMA) applied to human fMRI data. In contrast, to typical seed-based connectivity measures, FCMA comprehensively considers connectivity between every possible pair of voxels in the brain using sophisticated approaches to overcome historical computational limitations (Kumar et al., 2022; Turk-Browne, 2013; Wang et al., 2015). Specifically, FCMA uses a combination of machine learning (support vector machine; SVM) and parallel computing to efficiently map patterns of connectivity to stimulus or task information (Turk-Browne, 2013; Wang et al., 2015). The computational efficiency afforded by this approach is substantial in that it can reduce computation time from weeks to hours and has the potential to reveal information in fine-grained connectivity patterns that would be missed by conventional seed-based functional connectivity (Wang et al., 2015). We applied FCMA using an experiment that manipulated perception versus retrieval states while carefully controlling for three potentially confounding variables: (1) subjects performed the same judgments on the same images across both perception and retrieval tasks, minimizing differences in task demands and sensory content; (2) subjects were familiarized with the images in all conditions, minimizing differences in novelty/encoding across perception and retrieval; and (3) behavioral accuracy was matched across key conditions, minimizing differences in task difficulty.

To preview, we report four main findings: (1) background connectivity, which is orthogonal to evoked responses, allowed for classification of perception versus retrieval states with remarkably high accuracy; (2) perception and retrieval states can be discriminated parsimoniously based on background connectivity patterns in a relatively small number of clusters that span three functional communities: the control network, default mode network (DMN), and retrosplenial cortices (RSC); (3) connections within the control network were relatively stronger during perception states whereas connections within the DMN were relatively stronger during retrieval states; and (4) RSC shifted its coupling with the control network and DMN as a function of cognitive state, suggesting that it acts as a hub for transitioning between perception and retrieval.

## MATERIALS AND METHODS

### Subjects

Twenty-seven adults with normal or corrected-to-normal vision were recruited to participate for monetary compensation at Princeton University. Three subjects were excluded due to excessive head motion for a total of 24 subjects in the current sample (eleven reported male, mean age = 23.3 years). The Princeton University Institutional Review Board approved the study protocol, and all subjects provided informed consent. The sample size is on par with previous studies that examined functional connectivity changes with memory (Cooper & Ritchey, 2019) and used FC patterns to differentiate task states (Shirer et al., 2012).

### Materials

Stimuli consisted of 64 scene and 64 face images. The scene images were collected from the “Massive Memory” dataset (Konkle et al., 2010; http://konklab.fas.harvard.edu/#). The face images were obtained from the FEI face database (Thomaz & Giraldi, 2010; https://fei.edu.br/∼cet/facedatabase.html) and contained emotionally neural expressions. Scrambled images were generated as the weighted average between the actual image and its phase-scrambled version (Oppenheim & Lim, 1981; Stojanoski & Cusack, 2014). The script for creating the phase-scrambled images was adopted from code by Nicolaas Prins (e.g., as described here: https://github.com/rordenlab/spmScripts/blob/master/bmp_scramble.m).

### Experimental Design and Procedure

During the pre-scan training phase (∼20-30 minutes prior to scanning), subjects viewed randomly assigned pairs of images (always consisting of one face and one scene) and indicated how successfully they were in forming a mental association between the images. After viewing all stimuli pairs, they were given a 2-alternative force choice task (AFC) in which a face would be presented above two scenes (or vice versa) and the subjects had to indicate which of the bottom images had been paired with the top image before. The cycle of association-forming and 2-AFC task continued until subjects got each association correct two times in a row (**Figure 1A** **left**).

**Figure 1.**
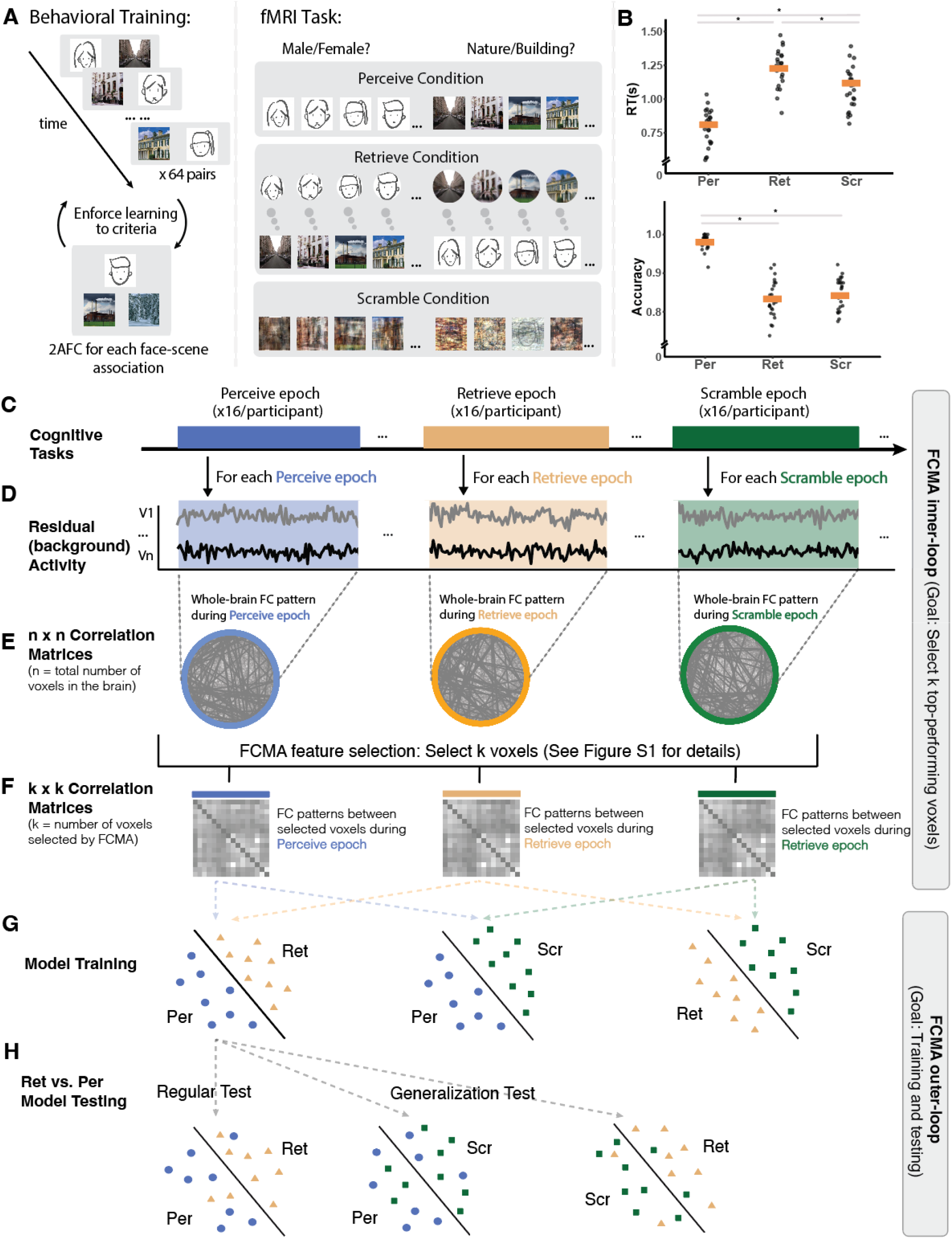
Task paradigm and analysis flowchart. (**A**) Behavioral training and in-scanner tasks. Subjects formed mental associations between face^1^ and scene images outside of the scanner and then made judgments on either perceptual or mnemonic information in the scanner. (**B**) In-scanner behavioral performance, with the orange bar indicating the sample mean and asterisks indicating *p* < 0.001. (**C**) Each condition had 16 epochs per subject, with each epoch lasting 40 s. (**D**) The residual activity for each task epoch was computed by regressing out the stimulus-evoked component using a finite impulse response general linear model. **(E**) Whole-brain voxel-wise background FC matrices were computed for each task epoch. These matrices were *n*-by-*n* shape, where *n* is the total number of voxels in the brain (*n* = 92,745). (**F**) Background FC matrices of FCMA-selected voxels. FCMA was implemented to select a certain number of voxels whose connectivity patterns were most accurate for separating perception and retrieval states based on two-way classifications of epochs from different conditions. The details of the feature selection process are shown in **Figure S1**. In essence, FCMA reduced the whole-brain correlation matrix to a *k*-by-*k* correlation matrix for each epoch, where *k* is the number of voxels selected by FCMA. The current study tested multiple values of *k*, ranging from 100 to 15,000. (**G**) Model training. Using a cross-validation framework, SVM classifiers with precomputed linear kernel were trained to separate all task condition comparisons. (**H**) Model testing. Trained models were tested using the left-out subject for each fold. During the regular cross-validation test, the model was tested on the same task condition comparison as it was trained on. During the generalization test, the model was tested on a different task condition comparison than it was trained on.

During the scanning session, subjects completed 3 task conditions across 6 functional runs (2 runs for each condition) using a block design. In the *Perceive* task condition, subjects were asked to identify the visual features of each cue on the screen. That is, if a face cue was presented, subjects were instructed to make a male/female judgment of the face via a button box, whereas if a scene cue was shown, subjects were instructed to make natural/man-made judgment of the scene. On the other hand, during the *Retrieve* task condition, subjects were asked to judge the gender or naturalness of the cue-associated image (i.e., the pair mate of the cue from the training phase). For example, if a face cue was shown, subjects needed to retrieve the specific associated scene image (not presented) and make a natural/man-made decision on that remembered information. Likewise, if a scene cue was shown, subjects were supposed to retrieve the specific associated face image and make a male/female decision on that remembered information. To ensure that the neural correlates we later identified were not driven solely by differences in task difficulty (as measured by accuracy) between *Perceive* and *Retrieve* task conditions, we included a *Scramble* condition. During the *Scramble* condition, subjects completed the same task as they did in the *Perceive* condition, but the visual cues were scrambled using a weighted average of the original image and its phase-scrambled version. The weight of this average was established based on a behavioral pilot study using the same design which sought to vary the weight until accuracy between the *Scramble* and *Retrieve* task conditions was matched (**Figure 1** **right**). Each run started with a 6-s blank lead-in period, followed by 8 task epochs and ended with a 6-s lead-out period. Each epoch consisted of a 4-s presentation of instructions, followed by 8 2-s presentations of single images presented at central fixation separated by a 1-s interstimulus interval. Each sequence of stimuli was followed by a 12-s inter-block-interval. Together, the duration of each trial, epoch, and run was 3 s, 40 s and 332 s, respectively. Note that all epochs within a functional run are of the same condition, and that all visual stimuli of an epoch are of the same category (i.e., either all faces or all scenes).

### Image Acquisition and Preprocessing

The fMRI data were acquired with a 3T scanner (Siemens Prisma) at the Princeton Neuroscience Institute. Functional data were acquired using a T2*-weighted multiband EPI sequence (repetition time = 1 s, echo time = 26 ms, flip angle = 50°, FOV = 260 x 260, resolution = 2.5 x 2.5 x 2.5 mm, multiband acceleration factor = 4) with 44 axial slices aligned to the anterior commissure/posterior commissure. A whole-brain T1-weighted MPRAGE 3D anatomical volume (1 x 1 x 1 mm voxels) was collected to improve registration. One phase and two magnitude field maps were collected to correct field inhomogeneities.

The first 6 lead-in volumes of each functional run were manually discarded before entering the preprocessing pipeline. Image preprocessing was performed using fMRIPrep 20.1.0rc1 (Esteban et al., 2019). All functional images were corrected for slice-acquisition time, head motion, and susceptibility distortion, and were normalized to a standard template, yielding preprocessed BOLD runs in MNI152NLin2009cAsym space. Following use of fMRIPrep, the minimally preprocessed functional runs were further processed using FSL (Woolrich et al., 2001) with a Nipype implementation (Gorgolewski et al., 2011). All functional images were smoothed with a 5.0 mm FWHM Gaussian kernel and high-pass filtered at 0.01 Hz. For each subject, the intensity values in each voxel in each of the 6 functional runs were then normalized using the mean and standard deviation of the resting period. The resting period for each functional run was defined as the 6 lead-out volumes plus all 12-s inter-block intervals of that run, which were shifted for 4 TRs to account for the hemodynamic delay. This normalization is intended to remove the BOLD signal differences across runs and thus all 6 runs were then concatenated to one time series and used for further modeling.

### Stimulus-Evoked and Residual (i.e., background) Activity

We computed residual activity by modeling and then regressing out the stimulus-evoked component from the preprocessed data in order mitigate stimulus-evoked coactivation confounds (Al-Aidroos et al., 2012; Cole et al., 2019). First, we constructed a confound regression model using FSL (implemented in NiPype) in order to minimize the effect of the following confound variables (obtained from fMRIPrep): six head motion parameters and the mean time series from white matter and cerebrospinal fluid. The resulting timeseries from regressing out the confound regressors are referred to as the stimulus-evoked activity timeseries in all subsequent analyses, as they are fully preprocessed, yet contain the stimulus-evoked components. Second, we estimated and removed the stimulus-driven components from the stimulus-evoked timeseries using a finite impulse response (FIR) model, which modeled the first 36 TRs for every epoch separately for face and scene epochs, resulting in 36 (TR) x 2 (epoch category) x 3 (condition) = 216 regressors. FIR is believed to be the optimal GLM for removing stimulus-evoked response because it does not assume the shape of the hemodynamic response function (Cole et al., 2019; Norman-Haignere et al., 2012). The residual timeseries data are referred to as the residual activity and used for computing background functional connectivity for all subsequent analyses.

### Full Correlation Matrix Analysis on Residual Activity

We utilized full correlation matrix analysis (FCMA) as implemented in the Brain Imaging Analysis Kit (BrainIAK; version 0.11; http://brainiak.org) to conduct an unbiased, whole-brain voxel-wise FC analysis that systematically considers all pairwise correlations in the brain to explore the differences in connectivity configurations between perception and retrieval states. All FCMA jobs were executed on Talapas, the HPC cluster at University of Oregon (https://racs.uoregon.edu/talapas). Each FCMA inner loop job was spread across 4 nodes, with each node supporting 28 threads and a job took around 4 hours to complete. Each node was equipped with two Intel E5-2690v4 processors, with 128 GB of memory. FCMA took in the residual activity and computed a full correlation matrix (i.e., whole-brain voxel-wise correlation matrix; 92745 voxels x 92745 voxels) for each task epoch. Therefore, for each subject, 8 (epoch) x 2 (run) = 16 full correlation matrices were computed per task condition (i.e., *Perceive*, *Retrieve* and *Scramble*; **Figure 1** **c-e**). Using these full correlation matrices, FCMA aimed to ( ) examine whether perception (*Perceive* and *Scramble*) versus retrieval (*Retrieve*) states can be successfully decoded from background connectivity patterns, and ( ) identify the connectivity configuration that characterizes each cognitive task state. Notably, the whole-brain correlation matrix is not easily interpretable by humans given its high dimensionality, and FCMA solves this problem by identifying of the most important/diagnostic regions of the brain involved in discriminating cognitive states (**Figure 1F**). Specifically, FCMA implements a nested leave-one-subject-out cross-validation (LOOCV) framework: the outer loop contains 23 training subject and 1 left-out test subject for each outer-loop iteration; and the inner-loop uses 22 training subjects and 1 left-out test subject within the outer training set for each inner-loop iteration. Importantly, the inner-loop intends to select the top *k* most useful voxels (based on their connectivity patterns) from the training data (**Figure S1**) and the outer loop aims to train classifiers on connectivity patterns of the selected voxels and test their ability to predict left-out data (**Figure 1G, H**). Note that we also performed parcel-level analysis to demonstrate the sensitivity advantage of our more fine-grained approach. We utilized the MNI version of the Schaefer parcellation scheme (https://github.com/ThomasYeoLab/CBIG/tree/master/stable_projects/brain_parcellation/Schaefer2018_LocalGlobal/Parcellations/MNI; Schaefer et al., 2018), and performed analyses on parcellations of different granularity (400 and 1000 parcels), which yielded comparable patterns of results as one another.

For each iteration of the outer loop, the inner-loop worked with the training data from 23 subjects (N-1 subjects) and performed a separate, nested LOOCV. Specifically, the inner-loop tested the accuracy of using each voxel’s seed maps (i.e., how much the voxel is connected to all other voxels in the brain) to differentiate perception (i.e., *Perceive* and *Scramble*) from retrieval (i.e., *Retrieve*) state (**Figure S1**). Each voxel would get an accuracy score for separating *Retrieve* and *Perceive* FC patterns, and another score for separating *Retrieve* and *Scramble* FC patterns. The minimum score of the two was assigned to the voxel and all assigned scores were averaged across the 23 inner-loop LOOCV iterations (i.e., training using 22 subjects and testing on a left out subject). The minimum score was used to make sure one of the two comparisons did not drive the overall effect and thus voxel selection prioritized discovery of regions which were sensitive to both comparisons. For example, a voxel’s seed map might be able to separate *Retrieve* from *Scramble* well by picking up the visual content information (i.e., scramble vs. intact), but not separate *Retrieve* from *Perceive* well when the visual content difference was absent. Computing the minimum accuracy scores allowed us to measure how well more comprehensively each voxel’s seed map encoded cognitive task state differences.

The resulting scores indicated the ability of each voxel to differentiate perception from retrieval states independent of the test data (i.e., the left out subject in the outer loop). Based on these scores, masks of the top *k* most useful voxels were created from each inner-loop and the whole-brain full correlation matrices were reduced to *k* x *k* correlation matrices. The current study tested all results with *k* = 100, 1000, 3000, 5000, 7000, 10000 and 15000 voxels, and subsequently focused on results using *k* = 3000. this was done for two reasons. First, model performances dramatically improved as the voxel masks were enlarged from *k* = 100 to *k* = 3000. However, model performance seemed to plateau when the top 3000 voxels were selected. Second, our information mapping pipeline (See **Methods: *Information Mapping***) identified a shared mask that included about 3500 voxels. The results from k = 3000 therefore provides the most direct comparison and is the most representative for subsequent analyses. Using these *k* x *k* connectivity matrices, the FCMA outer-loop trained 3 classifiers (one for each pair of task conditions, e.g., *Perceive* vs. *Retrieve*) and tested the model performance using the left-out testing data.

### Regular Cross-Validation and Generalization Tests

Classifiers built during FCMA were first trained and tested on the same task condition comparison (regular cross-validation). To quantify and compare model performance for each task comparison, we computed the area under the receiver operating characteristic curve (AUC) for each classifier. One way ANOVA was used to compare the AUCs for the three classifiers. The goals of the regular cross-validation tests were to examine o whether task conditions could be successfully decoded from background connectivity patterns and o whether task condition comparisons that involve cognitive task state differences (e.g., *Retrieve* vs. *Perceive*) could be decoded with higher accuracies relative to the comparison that did not involve such differences (e.g., *Scramble* vs. *Perceive*).

During the generalization test, a classifier was trained on one task condition comparison (e.g., *Perceive* vs. *Retrieve*) but tested on a different comparison (e.g., *Perceive* vs. *Scramble*). The generalization test can be conducted between each pair of task condition comparisons; we argue that the bidirectionally averaged generalization AUCs should indicate the classifiers’ sensitivity to a certain dimension of task differences (**Figure 2D**). For example, both the *Retrieve* vs. *Perceive* and *Retrieve* vs. *Scramble* comparisons involve cognitive task state differences. As a result, high bidirectionally averaged AUC scores across classifiers trained on *Retrieve* vs. *Perceive* and tested on *Retrieve* vs. *Scramble* and vice versa would indicate that the classifiers were indeed trained to pick up on cognitive task state information. On the other hand, high bidirectionally averaged AUC score across *Retrieve* vs. *Perceive* and *Scramble* vs. *Perceive* comparisons would suggest that the classifiers were trained to pick up on task difficulty differences. Likewise, high bidirectionally averaged AUC scores across *Scramble* vs. *Perceive* and *Scramble* vs. *Retrieve* comparisons would suggest that the classifiers were trained to detect scrambled versus intact visual features. Thus, the averaged bidirectional generalization accuracies served to measure the degree to which differences in cognitive task states, task accuracies, and visual content drove classifier performance. The primary goal of the generalization test, then, was to examine whether classifiers were able to preferentially detect connectivity patterns underlying perception versus retrieval task state differences.

### Multivoxel Pattern Classification Analyses

We trained activity pattern-based classifiers to test 1) whether FCMA classifier performances were driven by any coactivation confounds left in the residual activity and 2) whether simple MVPA and background FC patterns rely on the same type of cognitive processes. Per the first goal, using the *k* voxel masks generated by FCMA inner-loop (See section: ***Full Correlation Matrix Analysis on Residual Activity***), we trained pattern classifiers using residual activity to perform the regular cross-validation tests (**Figure S2**). Per the second goal, with each set of *k* voxel masks, we trained pattern classifiers using stimulus-evoked activity to perform the regular cross-validation test (**Figure 2B** **middle**). Pattern classification analyses were performed using a support vector classifier (C = 1) implemented in the Scikit-learn module in Python (Pedregosa et al., 2011).

### Information Mapping

Due to the leave-one-out cross-validation scheme, each fold of the outer-loop produced non-identical sets of top voxels. To obtain a common mask, we combined the FCMA inner-loop with non-parametric permutation tests to generate a group-level mask that included the top voxels differentiating perception from retrieval states better than chance. We obtained this group-level mask in three steps. First, using a LOOCV scheme across all 24 subjects, we averaged the classification accuracy composite score (i.e., the minimum of the accuracy scores for separating *Retrieve* vs. *Perceive* and *Retrieve* vs. *Scramble* FC patterns) for each voxel (**Figure S1**) and used this as the “observed” score for each voxel. Second, we shuffled the labels of all three task conditions for each subject and repeated the outer-loop training-testing pipeline using the shuffled labels for 100 iterations. As a result, we generated a null distribution of the classification accuracy composite scores for each voxel. Using the mean and standard deviation of the classification accuracy score null distribution for each voxel, we z-transformed the observed feature selection score computed above (i.e., the group-averaged composite accuracy score). We chose a relatively stringent voxel-wise primary threshold (P < 0.0001) to reduce the likelihood of false positives and to avoid finding large voxel clusters that span the brain (Woo et al., 2014). Voxels that passed the threshold were divided into clusters using 3dClusterize and the corresponding cluster-extent threshold was computed using 3dClustSim with AFNI (whole-volume alpha-values: -athr < 0.01). In total, 62 clusters passed the cluster-extent threshold, with size ranging from 620 to 2 voxels. In the last phase of the information mapping, we selected clusters according to their size and whether they improved model performance when combined with all larger clusters. Specifically, we began with the largest cluster (620 voxels), then we sequentially added voxels from the next largest cluster to our mask and used the FCMA outer loop to test how well this set of voxels differentiated perception from retrieval states. In other words, we iteratively ran a set of FCMA outer loop tests, and measured the change in classification performance as the mask accumulated each new cluster of voxels (**Figure S3A**). The rationale for performing cluster selection is to identify the smallest number of “sufficient” nodes whose network dynamics capture the differences between perception and retrieval states and ultimately reduce the full correlation matrix of the brain to a tractable set of representative ROIs.

### Community Detection

We applied community detection algorithms on the clusters identified by information mapping, aiming to understand the network level differences between perception and retrieval states. Specifically, we averaged across all voxels within each cluster to obtain cluster-level time series data for each epoch. Cluster-level background functional connectivity (FC) matrices were then computed for each epoch for every subject, and then averaged within each task condition, resulting in one group-averaged connectivity matrix for the *Perceive*, *Scramble,* and *Retrieve* conditions respectively. Community detection on weighted graphs (i.e., the connectivity matrices calculated above) was performed using the Louvain algorithm via NetworkX in Python. Following the approach used in prior work (Barnett et al., 2021; Ji et al., 2019), we ran the algorithm 1000 times on the weighted graphs to tune the resolution parameter in order to maximize modularity, which in turn captures how well a network can be subdivided into non-overlapping groups (Rubinov & Sporns, 2011). The tuned resolution parameters (gamma) were 1.10, 1.13, 1.19, for Perceive, Retrieve and Scramble task conditions, respectively. To visualize intra- and intramodular connectivity, we used the Fruchterman-Reingold force-directed projection implemented in NetworkX to project the graphs onto 2D spaces.

### Within- and Between-Communities Connectivity

To further understand network-level differences within and between the communities identified using the steps above (DMN, Control, and RSC; see **Results**), we computed a cluster-wise subject-level FC matrix for each task condition. We averaged the FC matrices for *Perceive* and *Scramble* conditions to create composite matrices that reflect FC dynamics underlying the perception state. To examine the coupling pattern across communities, we averaged across the connectivity strength between clusters (nodes) within the same functional community using both perception and retrieval state FC matrices. A paired sample t-test was used to examine whether DMN-Control connectivity strength differed between perception and retrieval states; and a two-way repeated measures ANOVA was then used to examine whether RSC nodes changed their coupling pattern with respect to the DMN and Control network nodes. To examine within-community connectivity strength, we averaged the connectivity strength between all nodes within the DMN, Control, and RSC regions for both perception and retrieval states. Similarly, a two-way repeated measures ANOVA was used to assess whether a functional community was biased toward a certain cognitive task state. All statistical analyses were performed using Pingouin 0.5.1 with Python3.

### Pattern Similarity Analyses on Stimulus-Evoked Activity

The goal of this set of analyses was to identify any potential differences between the three functional communities in their roles in performing cognitive tasks. We focused on three different cognitive aspects and tested the degree to which each functional community was sensitive to o the current visual category (i.e., face vs. scene), o the current behavioral task (i.e., gender vs. naturalness judgement), and/or o the current cognitive state (i.e., perception or retrieval). To do that, we performed pattern similarity analyses using both stimulus-evoked activity patterns and background connectivity patterns.

To obtain the stimulus-evoked activity patterns, we extracted the 24 task TRs for each epoch (after being shifted 4 s to account for hemodynamic delay) from the post-confound regression time-series. These stimulus-evoked estimates were then averaged along the time dimension and reshaped into a vector for each cluster (length of the vector is the number of voxels in that cluster). Background connectivity patterns for each cluster were computed as the correlation over time between the cluster and the other 15 clusters, reshaped into a 15-dimensional vector for each ROI (**Figure 6B**). To compute pattern similarity measures, the Fisher’s Z transformed correlations between each pair of vectors from different functional runs were calculated (Kriegeskorte et al., 2009; **Figure 6A**). Sensitivity was quantified as the difference between within-state epoch pattern similarity and between-state epoch pattern similarity. For example, when examining sensitivities for the current task judgement, pattern similarities were computed among all epochs with the same judgement (i.e., gender to gender and naturalness to naturalness) and compared to those with different judgments (i.e., gender to naturalness). Sensitivity was calculated by using the average within-state pattern similarity score minus the average between-states pattern similarity score. Thus, a significantly positive sensitivity index would suggest that the given ROI produced relatively distinct activity and/or connectivity patterns for the two aspects of the task (state, content, or judgement). We averaged the sensitivity scores across ROIs within the same functional communities. A one-way ANOVA was used to compare sensitivity scores of each cognitive process across the three functional communities.

## RESULTS

### Behavioral Results

We designed an fMRI task that required subjects to perform judgments on information that was either available in the perceptual environment (perception) or had to be retrieved from memory (retrieval), in both cases matched in terms of visual content and task difficulty (**Figure 1A**). Overall, subjects demonstrated greater accuracy in the *Perceive* compared to *Retrieve* condition, but comparable accuracy between *Scramble* and *Retrieve* conditions (**Figure 1B**). Specifically, one-way (condition) repeated measures ANOVAs revealed a main effect of condition for both reaction time (*F*_(2, 46)_ = 288.8, *p* < 0.001, *η*^2^ = 0.97) and accuracy (*F*_(2, 46)_ = 91.21, *p* < 0.001, *η*^2^ = 0.90). Although accuracy in the *Perceive* condition was significantly greater than that in the *Retrieve* condition (*t*_(23)_ = 13.4, *p* < 0.001, 95% CI = [0.16, 0.22], Cohen’s d = 3.55), the *Retrieve* and *Scramble* conditions did not significantly differ, serving as a useful point of comparison (*t*_(23)_ = -1.65, *p* = 0.11, 95% CI = [-0.07, 0.01], Cohen’s d = 0.45).

### Perception and Retrieval Involve Distinct Background Connectivity Patterns

We first examined whether perception and retrieval states involve different “state-related” whole-brain FC patterns. Using background connectivity analysis, voxel-wise whole-brain correlation matrix was computed for each epoch (**Figure 1E**; for details see **Methods**: ***Stimulus-Evoked and Residual Activity***). We then applied full correlation matrix analysis (FCMA) to test whether a trained support vector machine (SVM) could successfully separate perception epochs (*Perceive* and *Scramble*) from retrieval epochs (*Retrieve*; **Figure 1G-H**). To evaluate binary classifier performance within the top-performing voxels, we computed the receiver operating characteristic (ROC) curves and area under the ROC curve (AUC) for each classifier per subject, with larger AUC indicating better model performance (Hanley & McNeil, 1982). We started by having FCMA selecting *k* = 100 best performed voxels to train the classifier and gradually increased *k* until classifier performance asymptoted (for details see **Methods**: ***Full Correlation Matrix Analysis on Residual Activity***). As seen in **Figure 2A**, accuracy plateaued when the mask reached roughly 3,000 voxels, and the overall pattern of results across conditions did not dramatically change as a function of the mask size used. Accordingly, we performed follow-up analyses on the top *k* = 3,000 voxel masks. These analyses suggested that classifiers trained on background FC patterns successfully differentiated epochs of perception from retrieval states (*Perceive* vs. *Retrieve*: *M*_AUC_ = 0.87 ± 0.06, *_t_*_(23)_ = 28.62, *p* < .001, 95% CI = [0.84, 0.89], Cohen’s d = 5.84; *Retrieve* vs. *Scramble*: *M*_AUC_ = 0.83 ± 0.08; *t*_(23)_ 19.39, *p* < .001, 95% CI = [0.79, 0.86], Cohen’s d = 3.96; one-sample t-test against chance-level performance of μ = 0.5; see also **Figure S2A** for results in terms of proportion correct). Control analyses showed that state-related differences were selectively captured by background FC measures and not by left-over differences in the stimuli-evoked component of the signal. In particular, pattern analysis using residual activity patterns (i.e., simple MVPA on residual timeseries) in the same voxel masks failed to discriminate any task condition comparisons (all model performances ≈ 50% correct; **Figure S2B**).

One potential concern with the connectivity results above is that the above-chance classification performance was not strictly related to differences in perception versus retrieval states *per se* and might reflect the contribution of several potential confounding factors. For example, in addition to capturing differences in perception and retrieval, the *Perceive* vs. *Retrieve* comparison also varied in terms of task difficulty (i.e., accuracy) and the *Scramble vs. Retrieve* comparison also varied in terms of sensory input (i.e., partially scrambled vs. intact stimuli). To ensure that the variations in background FC patterns (among FCMA-selected voxels) were most strongly induced by state-related differences, we performed two extra sets of analyses to rule out these potential confounds. First, we tested the background FC pattern separability of *Perceive* vs. *Scramble* conditions—a task condition comparison that differed in terms of both task difficulty and visual content *but not* in perception/retrieval cognitive states. If the background FC patterns contain mostly state-related information, having equated perception state for both *Perceive* and *Scramble* should worsen classifier performance. Consistent with this hypothesis, AUC significantly differed between the non-state-related and state-related background FC classifiers (*F*_(2,46)_ = 20.30, *p* < 0.001, *η*^2^ = 0.47), with post-hoc tests revealing that *Perceive-Scramble* AUC was significantly lower than *Retrieve-Perceive* (*t*_(23)_ = -5.62, *p* < 0.001, 95% CI = [-0.19, -0.09], Cohen’s d = 1.45; **Figure 2A**) and *Retrieve-Scramble* (*t*_(23)_ = -4.10, *p* < 0.001, 95% CI = [-0.15, -0.05], Cohen’s d = 0.97). No significant difference was observed in the comparison of *Retrieve-Perceive* to *Retrieve-Scramble*, although there was a trend for higher AUC for *Retrieve-Perceive* (*t*_(23)_ = 1.82, *p* = 0.08, 95% CI = [-0.01, 0.08], Cohen’s d = 0.52).

In the second set of analyses, we leveraged classification generalization to assess the degree to which background FC patterns capture differences within each of three dimensions: (1) cognitive state (i.e., perception vs. retrieval); (2) task difficulty; and (3) visual content (for details see **Methods**: ***Regular Cross-Validation and Generalization Tests***). This was operationalized as how well a classifier trained on one pair of conditions generalized to another pair of conditions that differed on the same putative dimension (**Figure 1H**). We predicted that generalizability as measured by averaged bidirectional AUC should be greatest for task condition comparisons that involve state-related differences (i.e., *Perceive* vs. *Retrieve* to/from *Scramble* vs. *Retrieve*; **Figure 2B**) compared to task difficulty or visual content. Indeed, we found that background FC classifiers differed in their sensitivities to the three dimensions (cognitive task state: _AUC_ = 0.78 0.07; task difficulty: _AUC_ = 0.68 0.07; visual content: _AUC_ = 0.53 0.11; *F*_(2,69)_ = 52.80, *p* < 0.001, *η*^2^ = 0.60, **Figure 2C**). Follow-up t-tests suggested that generalization of background FC classification was highest when state-related differences were aligned compared to other dimensions of generalization (*ts* > 5.78, *p*s < 0.001). Together, these two analyses provide additional support to the finding above that the pattern of background FC contains information about perception and retrieval states above and beyond other differences between conditions.

**Figure 2.**
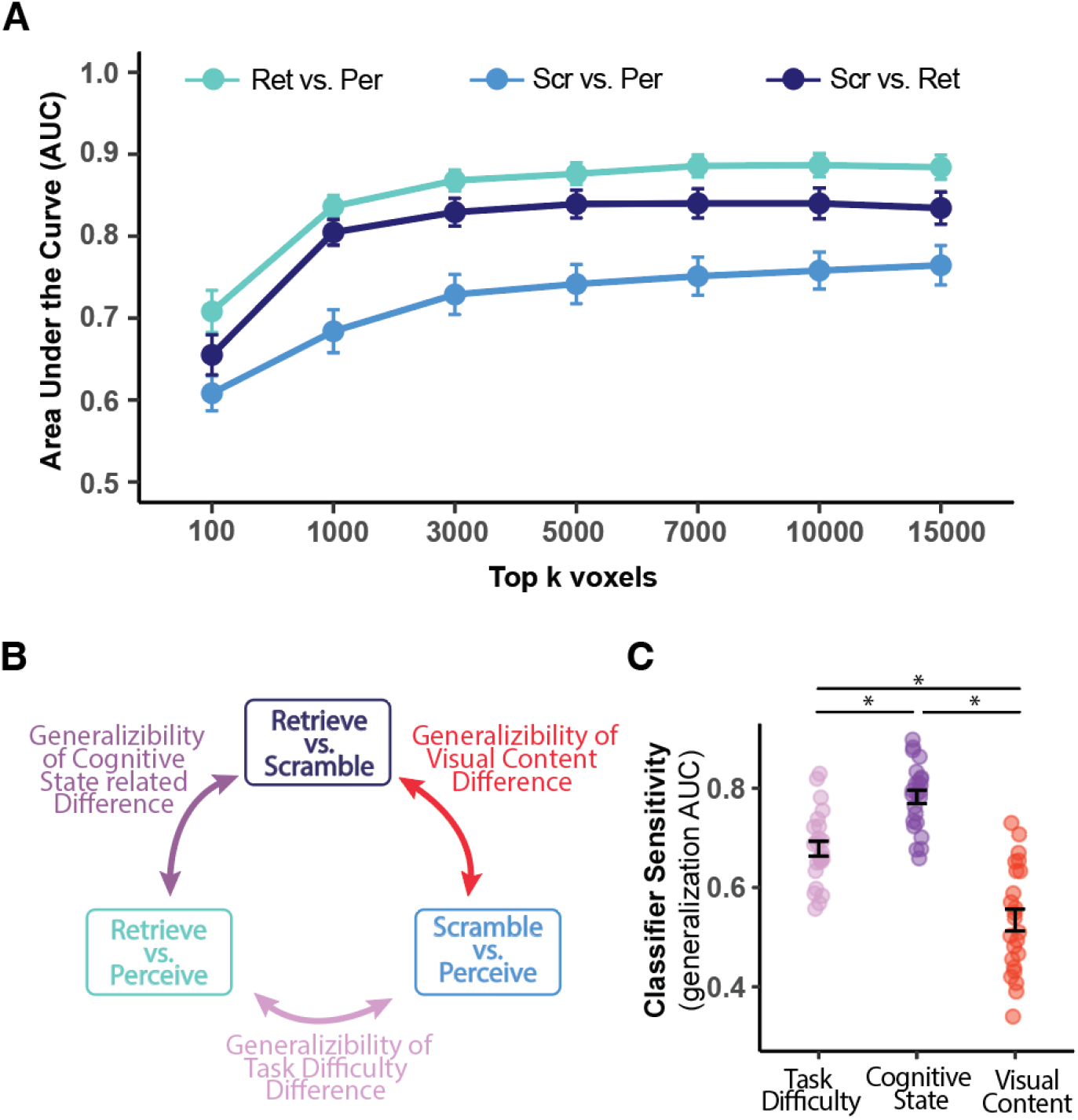
Classification of task conditions based on different features. **A**) Performance of background FC classifiers as a function of the number of voxels selected by FCMA. Area under the receiver operating characteristic curve (AUC) was computed for each leave-one-subject-out testing fold to examine the performance of the binary classifiers. The error bars indicate the standard error of the mean AUC across all subjects being tested. The results suggest that the top 3,000 voxels were able to differentiate task conditions as well as larger sets of voxels and so were used for follow-up analyses. **B**) Schematic diagram for measuring sensitivity to different aspects of cognitive processes using generalization tests. For example, given that both *Retrieve* vs. *Scramble* and *Retrieve* vs. *Perceive* comparisons involve state-related differences, success in classifying *Retrieve* vs. *Perceive* with the background FC classifier trained on *Retrieve* vs. *Scramble* (and vice versa) would provide strong evidence that the classifier is sensitive to state-related differences. **C)** Sensitivity of background FC classifiers to three distinctions: cognitive state (i.e., perception vs. retrieval), visual content (i.e., scrambled vs. intact), and task difficulty (i.e., low vs. high accuracy). Sensitivity was quantified as the average AUC from classifiers trained on one set of conditions and tested on another set along the same task-related dimension (generalization test). Error bars indicate the standard error of the mean across all testing folds. Asterisks indicate *p* < 0.05.

### Background Connectivity and Evoked Activity Patterns Capture Distinct Cognitive Processes

Previous research has shown that stimulus-evoked, multi-voxel activity patterns reflect ongoing cognitive processes and can be used to train classifiers for separating task conditions (Norman et al., 2006). How such measures relate to background FC is less well understood. Accordingly, we investigated whether patterns of stimulus-evoked activity reflect similar or distinct aspects of perception and retrieval states compared with patterns of background FC. Specifically, we trained MVPA classifiers on stimulus-evoked activity patterns for each pair of task conditions and compared the performance to classifiers trained using background FC patterns (**Figure 3** **left and middle**; for details see **Methods**: ***Pattern Similarity Analyses on Stimulus-Evoked Activity***). Stimulus-evoked activity patterns led to reliable classification of task conditions (*Retrieve* vs. *Perceive*: *M*_auc_ = 0.86 ± 0.06; *Scramble* vs. *Perceive*: *M*_auc_= 0.90 ± 0.12; *Scramble* vs. *Retrieve*: *M*_auc_= 0.90 ± 0.12; *ts* > 37.62, *p*s < 0.001). Interestingly, however, activity-based classification results demonstrated systematically different pattern compared to those of background FC-based classification. Specifically, a two-way repeated measures ANOVA with comparisons (*Retrieve* vs. *Perceive*, *Scramble* vs. *Perceive, Scramble* vs. *Retrieve*) and neural measure (stimulus-evoked activity, background FC) revealed a main effect of neural measure (*F*_(1,23)_ = 58.60, *p* < 0.001, *η*^2^= 0.72), highlighting the overall impact of using evoked-responses for classification. There was also a main effect of comparison (*F*_(2,46)_ = 16.19, *p* < 0.001, *η*^2^= 0.41). Strikingly, the ANOVA also revealed differential sensitivity across comparisons as a function of neural measure (*F*_(2,46)_ = 9.70, *p* < 0.001, *η*^2^ = 0.30). Based on post-hoc t-tests (see **Figure 3**), this interaction appeared to be driven by relatively greater classification performance for comparisons involving cognitive state differences (i.e., *Retrieve* vs. *Perceive* and *Scramble* vs. *Retrieve*) than those both involving perceptual decisions (i.e., *Scramble* vs *Perceive*), but only for background FC as the neural measure. Activity-based classification, on the other hand did not show a similar relationship.

The above findings are suggestive that stimulus-evoked activity and background FC may capture distinct aspects of the underlying cognitive state. To test this hypothesis more directly, we trained hybrid classifiers that combined both stimulus-evoked activity and background FC. The decision confidences of the hybrid classifier were computed as the averaged decision function outputs from both FC and MVPA classifiers. The rationale is that the hybrid classifiers should achieve better performance than either classifier on its own if the two neural measures capture distinct state-related processes (Manning et al., 2018). In line with the above results, a two-way repeated measures ANOVA revealed a significant main effect of classifier types (i.e., FC vs. MVPA vs. hybrid; *F*_(2,46)_ = 68.32, *p* < 0.001, *η*^2^ = 0.75; **Figure 3**). Follow-up analyses showed that the average AUC of hybrid classifiers was significantly greater than background FC classifiers (*t*_(23)_ = 9.95, *p* < 0.001, 95% CI = [0.11, 0.17], Cohen’s d = 2.58) and evoked-activity MVPA classifiers (*t*_(23)_ = 3.06, *p* = 0.006, 95% CI = [0.01, 0.04], Cohen’s d = 0.88). Together, these results suggest that evoked activity and background FC reflect distinct, possibly complementary, neural signatures of cognitive processes, with the latter displaying more relative sensitivity to state-related differences across retrieval and perception.

**Figure 3.**
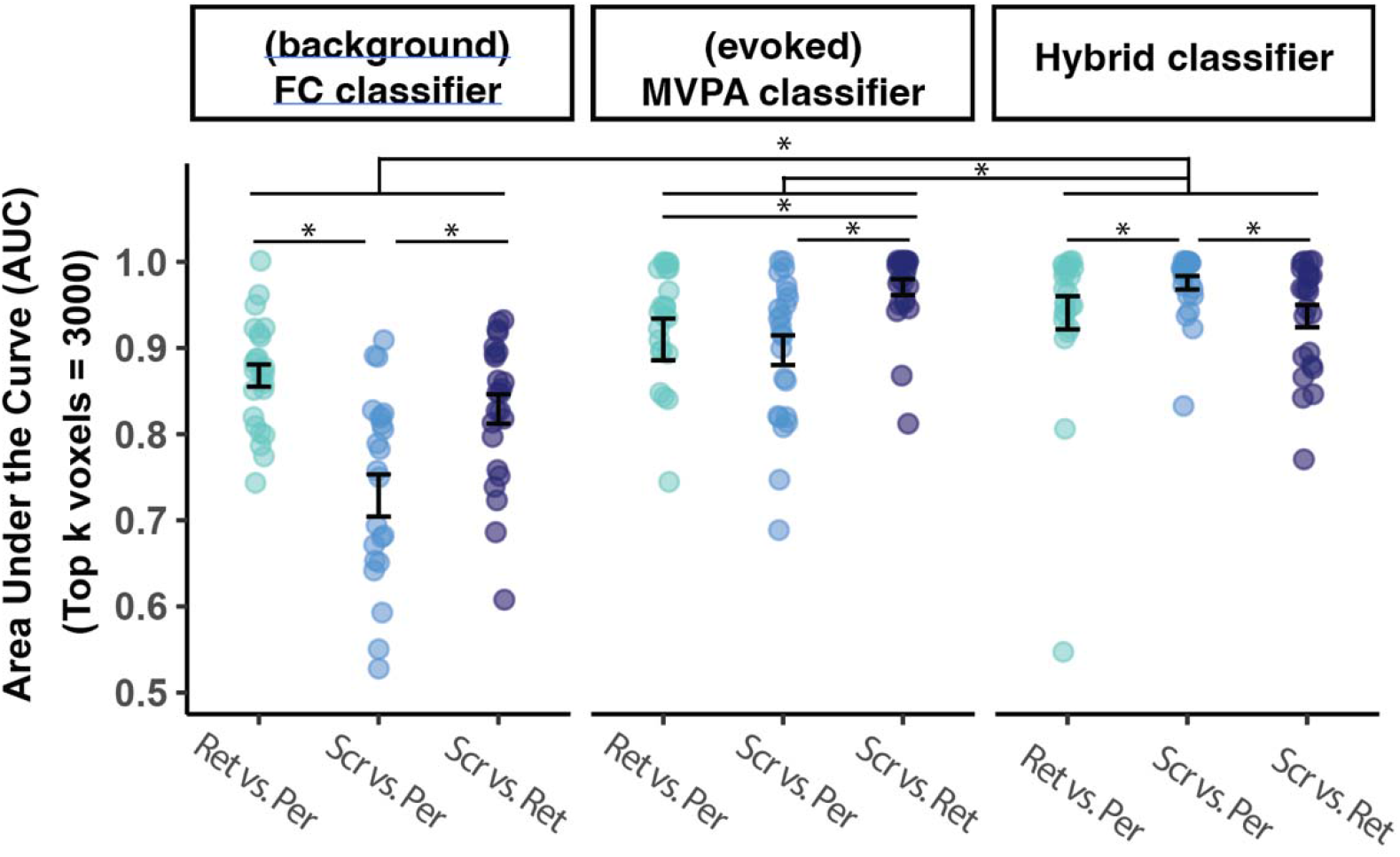
Classification of task conditions based on different neural measures. Comparing performance of binary classifiers trained with different neural measures: background FC (left), evoked activity (middle), hybrid FC + activity (right). Each dot represents a subject. The error bars indicate the standard error of the mean AUC across all subjects being tested.

### Regions and Functional Communities Underlying Perception and Retrieval States

The results so far suggest that background FC patterns of FCMA selected voxels capture differences between perception and retrieval states. However, the interpretation of these results in terms of brain regions is complicated by two key issues: 1) the top voxels were not necessarily identical across cross-validation folds and 2) the number of connections between voxels remains quite large. We sought to address each of these issues in turn. Per the first issue, because of leave-one-subject-out cross-validation, different sets of voxels were selected by FCMA for each left-out testing subject/fold. Accordingly, to allow further characterization and interpretation of state-related background FC patterns, we combined the FCMA inner-loop (**Figure S1**) with permutation-based statistical inference tests to obtain a shared set of voxels across all testing folds (yielding roughly 3500 voxels) whose background FC patterns captured state-related differences (for details see **Methods**: ***Information Mapping***). Per the second issue, even with a shared voxel mask, it may still be intractable to interpret FC patterns consisting of millions of connections (i.e., x 10^e^ unique connections among 3500 voxels). For this reason, we further reduced the dimensions of FC patterns using clustering. We first identified spatially contiguous clusters of voxels in the shared mask using a novel FCMA-then-clustering pipeline (see **Methods** and **Figure S3**). This process revealed 16 clusters of interests, whose cluster-level background FC patterns provided a parsimonious summary of the neural sources distinguishing between perception and retrieval states (**Figure 4A**; **Table S1**).

Notably, in addition to its greater interpretability, this FCMA-then-clustering approach also produced superior classification accuracy to the use of *a priori* parcellation-based clusters (Feilong et al., 2021; Schaefer et al., 2018). That is, by repeating the analysis pipeline with predefined brain parcels (instead of fine-grained voxel-wise analysis), the 16 clusters derived from voxel-level analyses had significantly greater discrimination performance compared to the best performing 16 predefined parcels in the parcel-level analysis (*F*_(1,23)_ = 96.71, *p* < 0.001, *η*^2^ = 0.81; **Figure S3B right**). This result held across different parcellation granularities (400 and 1000 parcels; Schaefer et al., 2018). Moreover, cluster-level background FC patterns retained their preference for state-related vs. non-state-related differences (**Figure S3**; *F*_(2,46)_ = 24.09, *p* < 0.001, *η*^2^ = 0.23).

Although the analyses above inform *which* brain regions might be most involved in differentiating perception and retrieval cognitive states, they do not provide information about *how* the regions are differentially connected. Indeed, previous research has suggested that functionally coupled brain regions form large-scale functional communities (Yeo et al., 2011) and that a cluster may be associated with different functional communities across different cognitive states (Braun et al., 2015). With this in mind, we next examined the functional community structures of the 16 clusters of interest during each task condition (i.e., *Perceive*, *Retrieve*, and *Scramble*). Specifically, we applied the Louvain community detection algorithm (Blondel et al., 2008) to the group-averaged background FC matrices for each condition (see **Methods: *Community Detection*** for details). Interestingly, the community structures underlying all three task conditions were consistent in that the 16 clusters were consistently partitioned into 3 functional communities (**Figure 4A**). The first functional community consisted of regions from the conventional default mode network (Buckner et al., 2008), including the bilateral inferior parietal lobule, precuneus, medial prefrontal cortex, posterior cingulate cortex, and the middle temporal gyrus, which hereafter we refer to as the DMN. The second functional community consisted of the bilateral prefrontal cortex, bilateral intraparietal sulcus, superior frontal gyrus, and temporal gyrus, most of which are part of the frontoparietal control network (Marek & Dosenbach, 2018); we refer to this as the Control network^2^. The last community consisted of ventral and dorsal retrosplenial cortices (RSC; Gilmore et al., 2016).

As an initial step to characterize the contribution of these different functional communities, we examined their evoked activation profiles during each task condition, using the averaged beta values estimated by an FIR model (**Figure S4**; see **Method: *Stimulus-Evoked and Residual Activity***). Consistent with network-level activity reported in past work, we found that DMN clusters deactivated (task-negative) during both perception and retrieval states, whereas Control network clusters activated (task-positive; Kim et al., 2015). Interestingly, only the activation profile of RSC clusters differed between perception and retrieval states: task-positive during the retrieval state but task-negative during the two conditions in the perception state.

**Figure 4.**
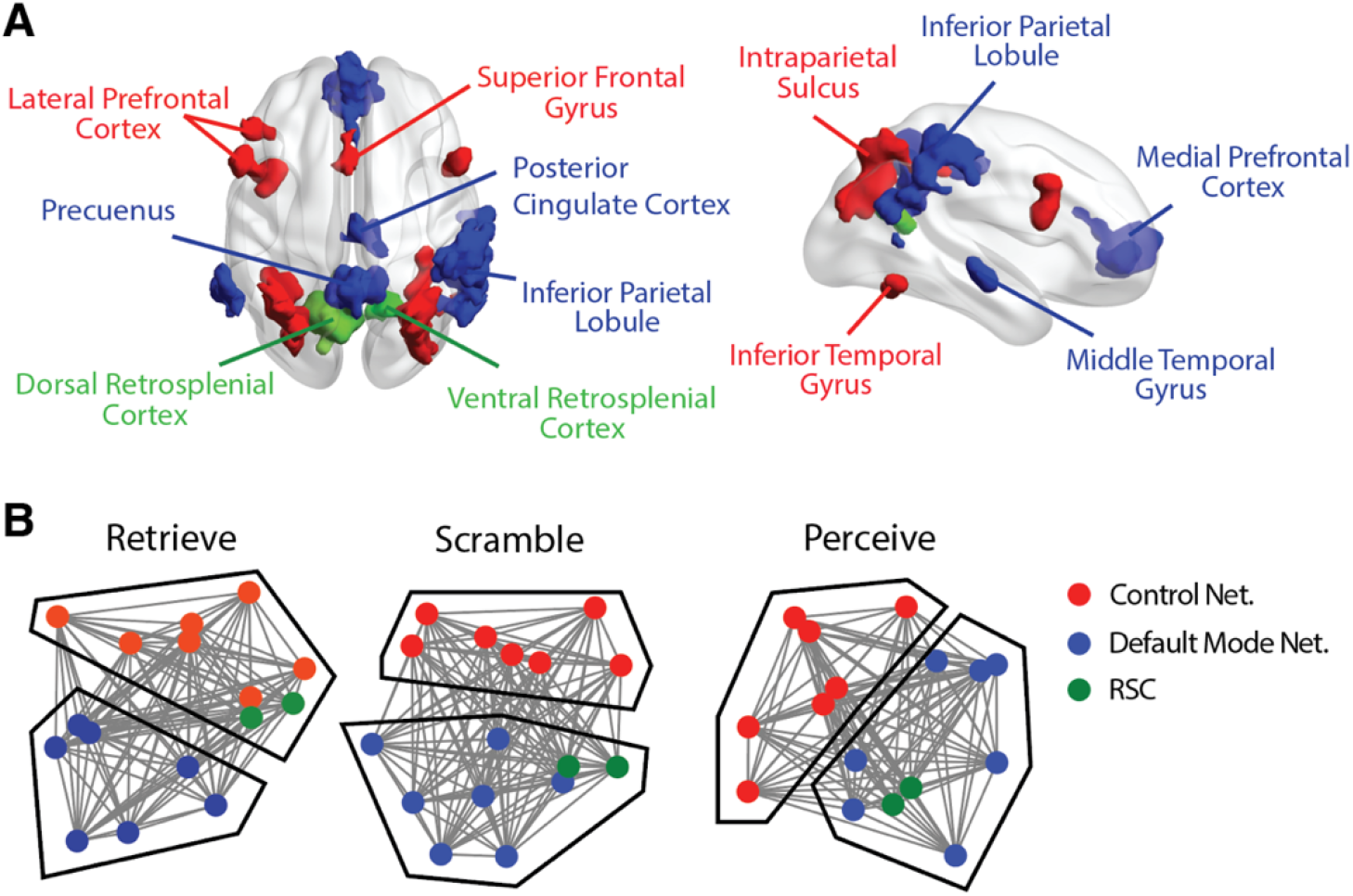
**A**) The 16 clusters of interest identified through the information mapping pipeline, partitioned into 3 functional communities. The color indicates the functional community assignment of each cluster. **B**) Force-directed graphs generated using the Fruchterman-Reingold algorithm implemented in NetworkX (V2.7.1; Fruchterman & Reingold, 1991). The projected physical distance in the graph indicates the degree to which nodes are being functionally connected. The color indicates the functional community allegiance of each cluster. Solid black lines represent manual delineations of the coupling pattern of RSC nodes.

### Within- and Between-Community Background FC Discriminates Perception from Retrieval States

Our FCMA-then-clustering pipeline successfully identified a tractable number of connections among selected brain clusters that robustly differentiated perception from retrieval states. In the next set of analyses, we aimed to characterize the nature of the difference in background FC by comparing within- and between-community FC strength across the two cognitive states. For each subject, we averaged across background FC matrices within each cognitive state (i.e., perception and retrieval). FC matrices from *Perceive* and *Scramble* conditions were thus averaged together in order to obtain a single measure of background FC for perception state for each subject. Note that we expected to observe background FC differences because the clusters were selected because of their sensitivity to state-related changes; the goal of this analysis is to interpret this difference. First, we sought to visualize the within- and between-community structure using force-directed plots which allow a concise representation of connectivity between all the regions. As can be seen in **Figure 4B**, connectivity between Control and DMN communities appeared to be stable across conditions, RSC was more closely connected with the Control network during retrieval (*Retrieve* condition) and with the DMN during perception (*Perceive* and *Scramble* conditions).

Next, in order to quantify within-community dynamics, we examined background FC for all cluster pairs within the same functional community during perception and retrieval states. **Figure 5A** shows the group-level differences in cluster-to-cluster background FC strength across the two states (perception minus retrieval). Visual inspection revealed that most intra-Control network connections (17 out of 21 connections) were stronger during the perception state, whereas the majority of intra-DMN connections (17 out of 21 connections) were stronger during the retrieval state. Quantitively, a repeated-measures ANOVA with factors of cognitive state (perception vs. retrieval state) and functional community (DMN, Control network, and RSC) revealed a significant interaction (*F*_(2, 46)_ = 19.05, *p* < 0.001, *η*^2^ = 0.45; **Figure 5B**). This interaction was driven by the fact that the averaged background FC strength among RSC clusters (*t*_(23)_ = 2.94, *p* = 0.007, 95% CI = [0.02, 0.1], Cohen’s d = 0.49) and cluster pairs within the Control network (*t*_(23)_ = 2.26, *p* = 0.03, 95% CI = [0.01, 0.06], Cohen’s d = 0.42) was stronger during perception compared to retrieval state (**Figure 5A** **left**), whereas it was numerically stronger among clusters in DMN during retrieval than perception state (*t*_(23)_ = 1.87, *p* = 0.07, 95% CI = [-0.06, 0.01], Cohen’s d = 0.32; **Figure 5A** **right**). Additionally, the ANOVA revealed a significant main effect of functional community (*F*_(2, 46)_ = 228.56, *p* < 0.001, *η*^2^ = 0.91). Clusters within the Control network had overall stronger connectivity density compared to those within in the DMN (*t*_(23)_ = 5.54, *p* < 0.001, 95% CI = [0.05, 0.1], Cohen’s d = 0.93). Note that this result holds even after accounting for the anatomical distances between clusters (**Figure S5**). Greater coupling between RSC regions may have possibly been driven by the fact that the two RSC clusters were close to each other anatomically. Lastly, the ANOVA did not show a significant main effect of cognitive state (*F*_(1, 23)_ = 2.06, *p* = 0.16, *η*^2^ = 0.08), suggesting that the overall background FC densities were comparable across perception and retrieval.

Finally, to better understand across network connectivity, we examined background FC for cluster pairs in different functional communities across perception and retrieval states. Specifically, as a function of the two states, we assessed each set of between-network connections separately, both in an individual, cluster-wise manner as well as averaged across all between-network connections. Based on the relative stability across conditions of Control/DMN communities compared to RSC seen in **Figure 4B**, we will present the results from Control/DMN connectivity and then connectivity with the RSC in turn. First, in terms of the Control/DMN communities, there were only hints of condition-dependent connectivity changes (**Figure 5C**). For example, rPFC (Control) tended to couple with DMN clusters more strongly during the retrieval state, whereas SFG (Control) had stronger coupling with DMN clusters during the perception state (**Figure 5D**). However, in aggregate, the averaged background FC strength of cluster pairs in Control network and DMN were comparable across the two cognitive states (*t*_(23)_ = 1.38, *p* = 0.18, 95% CI = [-0.1, -0.02], Cohen’s d = 0.39). In contrast, RSC shifted from coupling with the Control network to DMN, as cognitive states shifted from retrieval to perception respectively (**Figure 5E**). A repeated-measures ANOVA with factors of cognitive state (perception and retrieval) and functional community pair (i.e., the averaged connectivity measure between RSC nodes and regions in either Control or DMN regions) revealed a significant interaction (*F*_(1, 23)_ = 100.94, *p* < 0.0001, *η*^2^ = 0.81; **Figure 5F**). Specifically, RSC nodes had stronger averaged background connectivity with DMN nodes during the perception state (*t*_(23)_ = 4.83, *p* < 0.001, 95% CI = [0.05, 0.12], Cohen’s d = 0.76), but stronger background connectivity with Control nodes during the retrieval state. (*t*_(23)_ = 2.35, *p* = 0.03, 95% CI = [0.01, 0.09], Cohen’s d = 0.43). The ANOVA did not show a main effect of cognitive state (*F*_(1, 23)_ 1.22, *p* = 0.28, *η*^2^ = 0.05) or network pair (*F*_(1, 23)_ = 0.01, *p* = 0.91, < 0.01).

**Figure 5.**
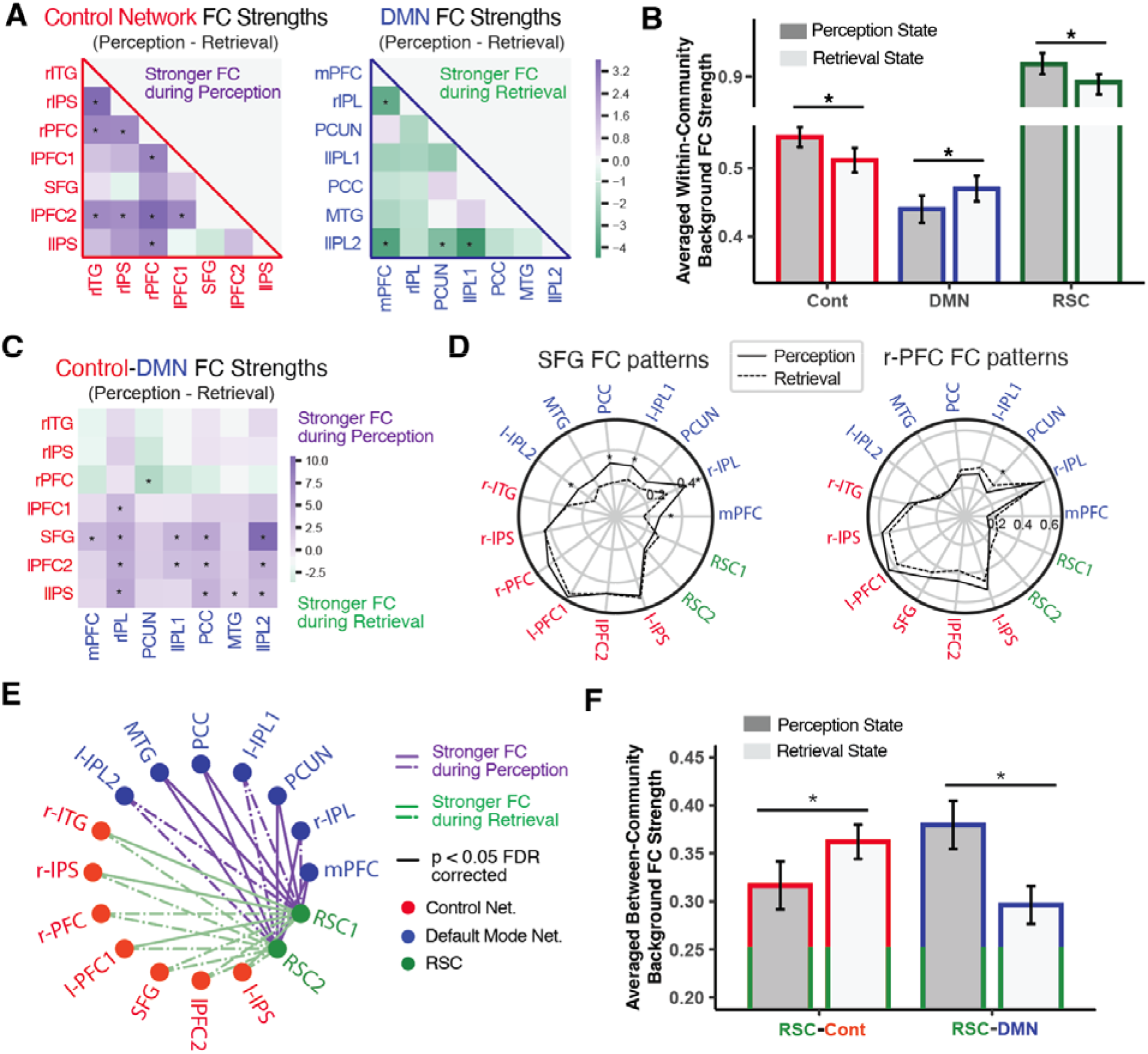
Connectivity configurations during perception and retrieval cognitive states. **A**) Group-level differences i background FC strength between each pair of clusters (within the same functional community) across perception and retrieval. The color of each cell represents the sign and magnitude of the *t* values, with a positive value indicating stronger coupling during perception state and a negative value indicating stronger coupling during retrieval state. Asterisks indicate *p* < 0.05 after FDR correction. **B**) Background FC averaged across all pairwise connections within the same functional community across perception and retrieval states. The error bars indicate the standard error of the mean across all subjects. Asterisks indicate *p*<0.05 **C**) Group-level differences in background FC between pairs of clusters in different functional communities (DMN and Control network) across perception and retrieval states. The color of each cell represents the sign and magnitude of the *t* values and asterisks indicate *p* < 0.05 after FDR correction. **D**) Background FC patterns of superior frontal gyrus (SFG) and right prefrontal cortex (rPFC) during perception and retrieval states. Asterisks imply *p* < 0.05 after FDR correction. **E**) Group-level differences i background FC between RSC clusters and DMN/Control clusters across perception and retrieval states. The color of each line represents the sign of the *t* values, with purple indicating stronger coupling during the perception state and green suggesting stronger coupling during the retrieval state. Solid lines indicate *p* < 0.05 after FDR correction. **F**) Background FC strength averaged across all pairwise connections between RSC clusters and DMN/Control clusters. The error bars indicate the standard error of the mean across all subjects. Asterisks indicate *p*<0.05

### Retrosplenial Cortex Plays Unique Role across Perception and Retrieval States

Although until now we have focused on differences between conditions related to cognitive state (i.e., perception vs. retrieval), performing the task required tracking two other forms of information: (1) the visual category^3^ currently being presented (i.e., face vs. scene) and (2) the behavioral judgement to perform (i.e., male/female vs. natural/manmade). Here, we performed pattern similarity analyses to assess how strongly these task components were represented in each cluster and functional community (**Figure 6A**). Further, given the divergence in sensitivity between background FC and evoked activity patterns reported above, we performed these analyses using both neural metrics. For each cluster, we calculated pattern similarity between pairs of epochs (**Figure 6B**) and compared within-class similarity (e.g., for visual category: face-face/scene-scene) and between-class similarity (e.g., face-scene) to index the cluster’s sensitivity to a given component. We then averaged across clusters within each functional community, yielding 3 sensitivity indices for each functional community per subject.

Pattern similarity measures of background FC suggested that clusters in different functional communities had different levels of sensitivity to cognitive state (**Figure 6C**; *F*_(2,46)_ = 6.32, *p* = 0.003, *η*^2^ = 0.22). In particular, RSC sensitivity was significantly greater than the Control network (*t*_(23)_ = 2.98, *p* = 0.006, 95% CI = [0.01, 0.05], Cohen’s d = 0.78) and numerically greater than the DMN (*_t_*_(23)_ = 1.95, *p* = 0.06, 95% CI = [-0.01, 0.04], Cohen’s d = 0.49). Although clusters were initially selected because their background FC patterns differentiated between cognitive states, perhaps leading to biased sensitivity estimates, this bias should not necessarily extend to differences in sensitivity between communities. Next, motivated by the differences between BG connectivity and evoked responses found above, we characterized pattern similarity using stimulus-evoked activity as well. Pattern similarity of evoked activity indicated that clusters in different functional communities also showed different level of sensitivity to visual category (**Figure 6D**; *F*_(2,46)_ = 48.35, *p* < 0.001, *η*^2^ = 0.68) and behavioral judgment (**Figure 6E**; *F*_(2,46)_ = 26.92, *p* < 0.001, *η*^2^ = 0.54). Further, the evoked activity patterns of RSC were significantly more sensitive to these two components than the DMN (visual-category: *t*_(23)_ = 9.02, *p* < 0.001, 95% CI = [0.12, 0.19], Cohen’s d = 2.26; behavioral-judgment: *t*_(23)_= 5.70, *p* < 0.001, 95% CI = [0.04, 0.08], Cohen’s d = 1.38) and the Control network (visual-category: *t*_(23)_= 4.52, *p* < 0.001, 95% CI = [0.05, 0.13], Cohen’s d = 1.21; behavioral-judgment: *t*_(23)_= 5.00, *p* < 0.001, 95% CI = [0.03, 0.07], Cohen’s d = 1.25). The one-way ANOVAs for all other communities across measures (activity or connectivity) failed to reach significance (*p*s > 0.25). Across metrics, these pattern similarity results highlight how RSC might be involved in many critical aspects of retrieving versus perceiving information (cognitive state, the visual content, and the behavioral judgment). However, interestingly, depending on whether the patterns are defined based on connectivity or activity, RSC can differentially capture the cognitive states of retrieval and perception relative to other regions.

**Figure 6.**
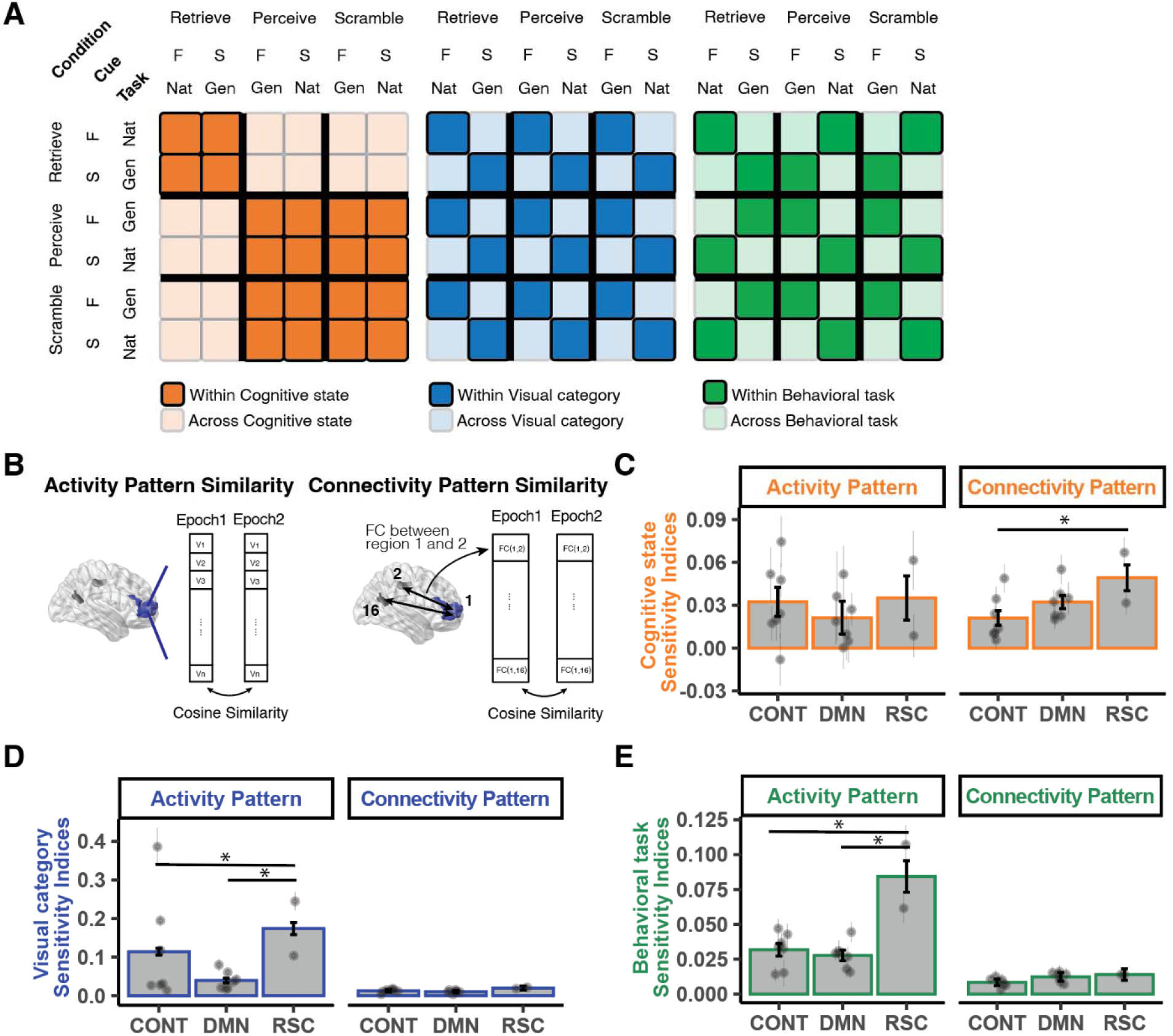
Pattern similarity analyses using both background FC patterns and stimulus-evoked activity patterns. **A**) Diagram for computing pattern similarity measures between pairs of epochs for each of the three task components (cognitive state, visual category, behavioral task). Darker colors indicate epochs of the same class (e.g., visual categories being face-face) whereas light colors indicate epochs of different classes (e.g., visual categories being face-scene). **B**) Diagram for computing pattern similarity measures with both FC and activity patterns. When computing activity pattern similarity measures, each epoch is represented by a *n*-dimensional vector, where *n* is the total number of voxels in the respective cluster. When computing connectivity pattern similarity, each epoch is represented by a 15-dimensional vector, representing the background FC measure between this cluster and the other 15 clusters. **C**-**E**) Sensitivity indices of each functional community with regard to each of the three task component. Sensitivity indices were quantified as the average difference of within- vs. between-class epoch similarity. Each individual dot indicates the average across all subjects for a cluster and the error bars indicate the standard error of the mean across all subjects. Each bar represents the average across all subjects for a functional community and the error bars indicate the standard error of the mean across all clusters within the respective functional community. Asterisks indicate *p* < 0.05.

## DISCUSSION

The goal of the current study was to characterize and differentiate perception versus retrieval states in a way that captures the complexity of whole-brain functional connectivity (FC). First, we found that patterns of background FC across perception versus retrieval states were systematically different from one another (**Figure 2A**). Moreover, the differences were best captured by background FC patterns between 16 clusters across 3 hypothesized functional communities (**Figure 4**; **Table S1**). Our whole-brain analysis pipeline allowed us to extend findings from previous research (Cooper & Ritchey, 2019) by identifying important brain clusters and coupling patterns beyond memory-related brain regions (**Figure 5**). Second, our results showed that background FC and evoked activity tend to capture distinct component processes (**Figure 3**), with the former being more “state-related” and the latter being more “stimulus-related” (Summerfield et al., 2006). Third, we demonstrated the utility of full correlation matrix analysis (FCMA; Kumar et al., 2022; Wang et al., 2015) and showed how the feature selection process of FCMA, paired with cluster-based dimensionality reduction, can be used to improve the interpretability of high-dimensional FC results (**Figure S1, S3**). We conclude by highlighting how the above findings are consistent with a framework of selective attention where, given the same perceptual input, directing attention externally would promote perception of the input-related sensory features, whereas directing attention internally would promote retrieval of the input-associated episodes (Chun et al., 2011).

### Background Functional Connectivity Configurations Underlying Perception and Retrieval States

The current study had subjects use the same visual input as either the target of a perceptual judgement or the trigger for episodic memory. Our results suggest that in order to successfully perform tasks in these different conditions the brain produces distinct background FC configurations to maintain a sustained cognitive state for either perception or retrieval (**Figure 2A**). Similar state-related shifts in background FC configurations have been reported in previous work. Cooper and Ritchey (2019) found that pre-defined regions of interest in memory-related brain systems (anterior temporal and posterior medial networks; Ranganath & Ritchey, 2012) showed stronger background FC during retrieval over perception. Using a purely data-driven approach, a subset of our findings are largely consistent with Cooper and Ritchey (2019), as we found that the retrieval (vs. perception) state is characterized by stronger background coupling between clusters in the default mode network (DMN), which is important for internal cognitive processes (Buckner et al., 2008; Yeshurun et al., 2021). Specifically, our pipeline highlighted seven clusters in the DMN (blue clusters in **Figure 4A**; **Figure 5A, B**), including the bilateral inferior parietal lobule (IPL), posterior cingulate cortex (PCC), precuneus (PCUN), medial prefrontal cortex (mPFC) and middle temporal gyrus (MTG). Note that IPL, PCUN, and PCC are considered parts of the posterior medial (PM) network for memory-guided behaviors (Ranganath & Ritchey, 2012), and IPL, PCC and mPFC have been shown to form a “core recollection network”, supporting memory retrieval success (Rugg & Vilberg, 2013). Indeed, several studies have showed that functional interactions between these regions contribute to different aspects of episodic memory (Cooper & Ritchey, 2019; Geib et al., 2017; King et al., 2015). In particular, the interaction between the IPL and PCUN may connect episodic features to form an integrated neural representation, while conceptual knowledge and existing schemas are integrated by PCC and mPFC (Ranganath & Ritchey, 2012; Ritchey & Cooper, 2020).

Our findings further extend previous work by showing that the perception (vs. retrieval) state is characterized by increased background FC within a functional community with clusters in both the Control Network and the Dorsal Attention network (red clusters in **Figure 4A**; **Figure 5A, B**). Although there are situations where these networks might be engaged during memory-related processes (Hutchinson et al., 2014; Rosen et al., 2016, 2018), much evidence suggests their consistent involvement in processing the external world. That is, previous research has consistently implicated these regions as being a part of a larger task-positive network that is typically engaged when the brain processes external stimuli (Fornito et al., 2012; Golland et al., 2008). Together with the results concerning the DMN, our findings are consistent with the idea that the brain has two anatomically separable systems that primarily correspond to “externally oriented” versus “internally oriented” processing (Golland et al., 2008). Although our analyses did not implicate the hippocampus (see below), an interesting possibility is that the hippocampus may switch ‘modes’ of processing (externally- vs. internally-oriented) and consequently drive the network-level differences observed in cortex here (Duncan et al., 2012; Hasselmo et al., 1996; Honey et al., 2017).

Our whole-brain data-driven approach also revealed the importance of the background FC patterns in retrosplenial cortex (RSC; green clusters in **Figure 4A**) for accurately characterizing perception versus retrieval states. Specifically, we showed that RSC-DMN coupling was substantially greater during perception whereas RSC-Control coupling was greater during retrieval (**Figure 5E**, **F**). This connectivity pattern may seem counterintuitive at first, but interestingly it replicates findings of previous research. RSC is hypothesized to be part of the PM network, and its background FC patterns were also examined in Cooper and Ritchey (2019). Their results suggested that, although most ROIs in the PM network showed stronger coupling with each other during retrieval compared to perception, the background FC patterns of RSC did not demonstrate observable enhancement during retrieval state with any other regions in the PM network (c.f., Figure 3C right in Cooper & Ritchey, 2019). Instead of showing stronger coupling with PM regions during the retrieval state, they found that RSC had numerically stronger background FC with task-positive regions, such as the inferior temporal cortex, which is consistent with what we observed (**Figure 5E**). Benefiting from the whole-brain data-driven approach, the current study revealed a more complete picture of RSC background FC during perception and retrieval states, identifying regions in the DMN and Control networks beyond classical memory-related systems (Cooper & Ritchey, 2019).

Although the findings in RSC are consistent with past observations, conceptual interpretation of its role here remains speculative. We do note that previous human fMRI and rodent studies suggest that RSC is involved in connecting external and internal states (Bicanski & Burgess, 2018; Yeshurun et al., 2021). For example, a study in mice found that RSC integrates both allocentric mapping (the animal’s location in the external world) and egocentric frame (the animal’s internal representation of the location) to navigate through a maze (Alexander & Nitz, 2015) by combining sensory inputs and mnemonic information from the medial temporal network (Bicanski & Burgess, 2018). Similarly, human RSC has been proposed to be a hub for connecting external and internal worlds (Yeshurun et al., 2021), such that it integrates external cues with self-generated information to guide behavior (Ranganath & Ritchey, 2012). In this respect, background FC in RSC during perception and retrieval may capture the role of RSC in bridging perceptual and mnemonic information; however, it is unclear why RSC would express higher coupling with the functional community putatively less involved with the task at hand (e.g., with DMN during the perception state). Accordingly, we believe an important direction of future research will be to fully characterize how the RSC assists in establishing perception and retrieval states.

Whereas some prior studies have identified changes in background connectivity within the hippocampus (Bein et al., 2020; Duncan et al., 2012; Honey et al., 2017) the hippocampus did not pop up using our whole brain approach. In particular, these studies have found that coupling between CA1 and CA3/DG is stronger during retrieval compared to perception states. Moreover, computational models also suggested that neuromodulations in the hippocampus (e.g., Acetylcholine shifts) can be crucial in switching between perception and retrieval states (Hasselmo & Schnell, 1994; Tarder-Stoll et al., 2020). By contrast, clusters identified in the current study did not highlight the importance of hippocampal subregions. We believe that this could be due to the fact that FCMA was configured to generate group-level voxel masks, ignoring individual differences in hippocampal subfield anatomy. The hippocampus is a relatively small brain region, and its subfields often include on the order of dozens of voxels (under typical scanning parameters) that may differ in their precise anatomical location across individuals. As a result, analyses of hippocampal subregions require subject-level hippocampal subfield segmentation, with each subject obtaining a slightly different segmentation map. In this respect, a shared group-level voxel mask may not be able to capture the important but subtle interactions among hippocampal subfields.

### Background FC Captures “State-Related” Signals and Evoked Activity Reflect “Stimulus-Related” Signals

Different mental states can be reflected in distributed and overlapping patterns of evoked activity in the brain. An influential line of work has aimed to decode this information using the technique referred to as multivariate pattern analysis (MVPA; <u>Norman et al., 2006)</u>. At the same time, another line of complementary work has investigated the inter-regional connectivity structure of the brain by examining the patterns of BOLD correlations (functional connectivity; FC) between multiple brain regions (Smith, 2012). Both approaches have been fruitful, resulting in tremendous insights into our understandings of human cognitive processes (Haxby, 2012; Song & Rosenberg, 2021). Importantly, some previous research has suggested that these two neural measures are likely to capture and reflect distinct, or at least non-overlapping aspects of cognitive processes. For example, Song et al. (2021) found that when viewing or listening to narratives, ongoing attentional engagement can only be predicted by FC-based models, whereas models trained with regional activity patterns failed to capture this information.

Additionally, Manning et al. (2018) showed that an ensemble model that relied on both FC- and activity-based neural measures outperformed models that utilized either measure on its own. This result suggests that FC and activity patterns can capture partially non-overlapping variance in cognitive processes.

Extending these findings, the current study suggests that background FC-based measures are more sensitive to differences in cognitive states whereas activity-based measures are more likely reflect differences in stimulus-related features of task. Specifically, we found that background FC-based classifiers better separated task conditions that involved state-related (i.e., perception vs. retrieval) than those only involved stimulus-related differences (e.g., visual content). On the contrary, MVPA classifiers did not demonstrate a clear preference for state-related comparisons over stimulus-related distinctions (**Figure 3**). Our findings are in line with the theory that the fMRI data acquired at each voxel are composed of both state-related and stimulus-related activity, and that FC-based measures could be better suited to capture “state-related” signals whereas activity patterns better capture the “event-related” component (Summerfield et al., 2006). It is also worth noting that there are different types of FC-based measures; the current study primarily tested the background FC measures by regressing out the stimulus-evoked component using a general linear model (Al-Aidroos et al., 2012; Bejjanki et al., 2017; Norman-Haignere et al., 2012; Tompary et al., 2018). However, some studies computed their FC measures without regressing out the stimulus-evoked component (e.g., Song et al., 2021) or rely on stimulus-evoked parametric measures (e.g., beta series correlation; Bein et al., 2020). Future work might further examine whether these types of FC-based measures also preferentially capture state-related aspects of the cognitive process.

### The Utility of Feature Selection in Whole-brain Voxel-wise FC Analyses

The current study used full correlation matrix analysis (FCMA; Kumar et al., 2022; Wang et al., 2015) to explore whether cognitive states are encoded in whole-brain voxel-wise background FC patterns. Our approach is systematically different from those used in previous studies, such as connectome-based predictive modeling (Shen et al., 2017) in two ways. First, FCMA operates on voxel-level connectivity matrices instead of the commonly-used, lower-dimensional parcel-averaged time series. Second, FCMA uses a nested leave-one-subject-out cross validation framework that enables more efficient feature selection to reduce large connectivity matrix to a tractable size. Here we discuss the potential trade-off regarding these two distinctions. First, our results suggest that FCMA identified regions that, even after clustering over contiguous voxels, retained a higher sensitivity for differences in cognitive states compared to parcel-level analyses (**Figure S3B right**). This finding is consistent with previous research suggesting that voxel-/vertex-level FC patterns are more sensitive to other cognitive measures (concerning intelligence) compared to relatively coarse-grained parcel-level analysis (Feilong et al., 2021). The differences in sensitivity could be due to the fact that many parcellation schemes were defined from whole-brain resting-state functional connectivity profiles rather than task-based connectivity and previous work has suggested that the functional architecture of the brain might change across resting and task states (Cole et al., 2014) and may also vary across different task states (Krienen et al., 2014). It is worth mentioning that the use of predefined parcellation schemes can significantly improve sensitivity (as measured by effect size) in other types of analyses (e.g., univariate analyses) compared to voxel-/vertex-level analysis (Li et al., 2021). Thus, future studies need to further investigate the tradeoffs for predefined parcellation schemes in connectivity analyses.

The second way the current approach differs from prior approaches is in terms of interpretability. That is, machine learning models often face a trade-off between prediction accuracy and model interpretability (Feilong et al., 2021). In the context of FC-based models, previous work has primarily focused on constructing a model for making the most accurate predictions on trait-like demographic variation (Finn et al., 2015), behavioral performance (Rosenberg et al., 2016, 2020), or task conditions (Gonzalez-Castillo et al., 2015; Shirer et al., 2012). Despite high prediction accuracies in these models, it is often hard to interpret the FC configuration given the vast number of connections. For example, Rosenberg et al. (2016) identified a “high-attention network”, consisted of 757 edges across the entire brain, whose pattern of connectivity reliably predicted better performance on a sustained attention task. Indeed, such an approach likely captures many nuances in the neuronal distributed process of attention and has high predictive accuracy. However, it is possible that such high accuracy comes at the expense of interpretability. That is, constraining the number of edges and nodes in the descriptive network might sacrifice some degree of prediction accuracy, but the resulting FC configuration might be more easily interpreted. The current study attempted to do this by combining a whole-brain voxel-level FC model with feature selection using a nested cross-validation framework. Specifically, we quantified the “utility” of each connection using the machine learning training and testing framework (See **Method**: ***Full Correlation Matrix Analysis on Residual Activity***; **Figure S1**). As a result, we were able to select the most useful connections in an unbiased, automated fashion, reducing large correlation matrices to a tractable size.

Remarkably, the model based on connections among only a set of 16 regions retained comparable AUC scores compared to models based on connections among 3000 voxels (Figure 2a; 0.83 vs. 0.87)^4^. Thus, the current study demonstrates the value of using the FCMA voxel-to-cluster pipeline in order to yield the most relevant and interpretable FC configuration profile while largely maintaining prediction performance.

### Ideas and Speculation: Perception Versus Retrieval as Outcomes of Selective Attention

Reconfiguration of background FC patterns has been examined as a neural mechanism underlying, or resulting from, selective attention (Desimone & Duncan, 1995). Specifically, the “switching-train-track” framework for selective attention argues that selective attention prioritizes goal-relevant information by strengthening the interaction (as measured by the changes in background FC) between goal-relevant brain regions (Al-Aidroos et al., 2012; Miller & Cohen, 2001; Turk-Browne, 2013). For example, previous work has shown that, when presented with face-scene composite stimuli, early visual cortex shows stronger intrinsic FC coupling with face-specialized regions (e.g., fusiform face area) when subjects attended to the face component but with scene-specialized regions (e.g., parahippocampal place area) when subjects attended to the scene component (Al-Aidroos et al., 2012; Córdova et al., 2016; Norman-Haignere et al., 2012; Tompary et al., 2018). Here we speculate that the difference between perception and retrieval could be understood and investigated as an key dimension of selective attention. Specifically, a recent taxonomy dichotomized attention into external attention, with the target being perceptual information and internal attention, with the target being self-generated trains of thoughts (Chun et al., 2011). Under this dichotomy, perception versus retrieval can be thought of as a form of external vs. internal attention states for information processing. When a given input is being attended externally, the sensory information of the input is being preferentially processed while the mnemonic information is suppressed. On the other hand, when the same input cues internally-directed attention, the mnemonic information will be enhanced while the sensory information is suppressed (Chun et al., 2011; Gazzaley & Nobre, 2012). The switching-train-track account of selective attention has been mostly studied by examining FC changes within the ventral visual stream when subjects were attending externally to different visual categories (e.g., face vs. scene). The current study has provided evidence that this account of selective attention may generalize to the whole brain and to more complex cognitive states (e.g., perception vs. retrieval). In particular, although previous research studying perception vs. retrieval has focused primarily on decoding the “train of thoughts” (i.e., stimulus-related features; Rissman & Wagner, 2012), the selective attention perspective highlights the importance of investigating state-specific “train tracks”—i.e., background FC patterns—and understanding how such train tracks establish and sustain particular cognitive states.

## Data and Code Availability

Processed fMRI data including both the stimulus-evoked and residual time series supporting the primary findings of this study are available on the Open Science Framework (OSF) at https://osf.io/yfwc7/. Scripts for performing and reproducing the specific analyses described in this paper can be found through Github at https://github.com/peetal/Decode_AttenStates.

## Funding and Acknowledgement

This work was supported by the Intel Corporation, the John Templeton Foundation, and NIH R01 EY021755. We express our deepest thanks for the feedback we received from Dr. Ken Norman and Dr. Sam Nastase. We are also grateful for helpful discussions and support from members of the Hutchinson Lab of Cognitive Neuroscience, the Dubrow Lab, and the Kuhl Lab.

---

**Supplementary Figure 1.**
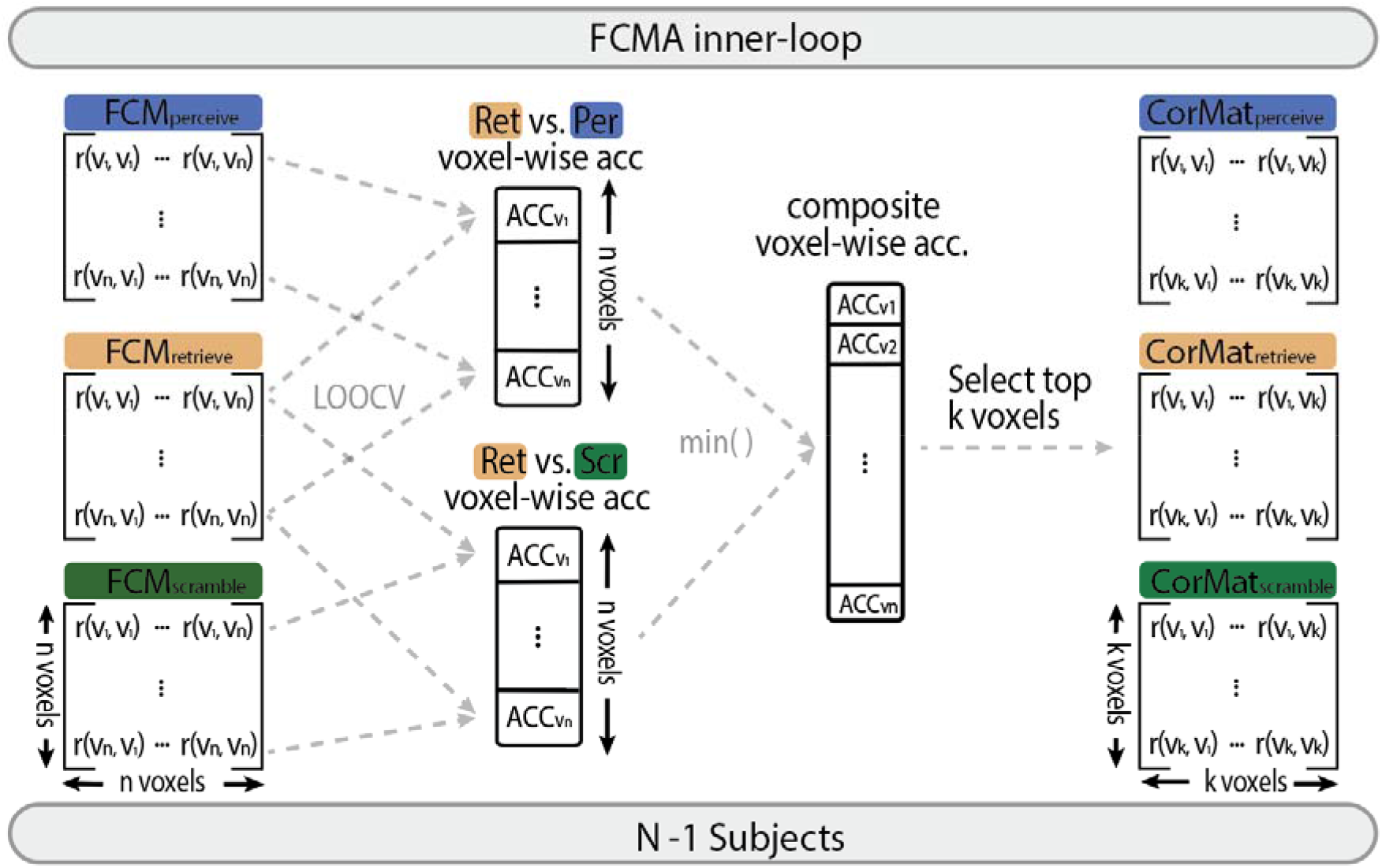
FCMA feature selection process. The FCMA inner-loop used a leave-one-subject-out cross validation (LOOCV) framework to quantify the utility of each voxel. Within each training fold, we examined the degree to which each voxel’s background functional connectivity patterns (seed map) can be used to differentiate *Retrieve* vs. *Perceive* conditions and *Retrieve* vs. *Scramble* conditions. This leads to two *n*-vector accuracy measures across voxels, one for each task condition comparison. A composite voxel-wise accuracy score was computed by taking the minimum of the two accuracy values for each voxel. The top *k* voxels in terms of composite accuracy score were selected to construct dimensionality reduced FC patterns (Figure 1F). Because each fold’s training data differed by one subject, a different (but typically highly overlapping) set of *k* voxels could be selected for each training fold. As a result, the full cross-validation framework create 24 masks of selected voxels for each choice of *k*.

**Supplementary Figure 2.**
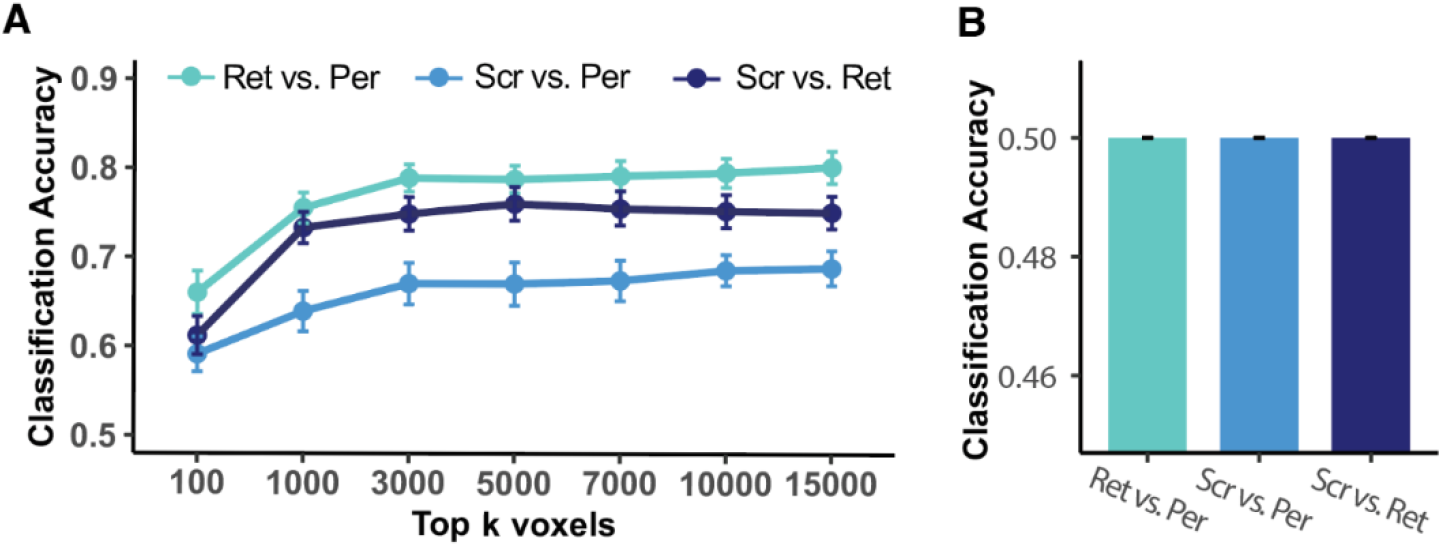
**A)** Background FC classification accuracy for each task comparison across different numbers of voxels selected by FCMA. Performance asymptoted when *k* = 3,000: *Perceive-Scramble*, *M*_acc_ = 67% 11%; *Perceive*-*Retrieve*, *M*_acc_ = 79% 8%; *Retrieve*-*Scramble*, *M*_acc_ = 75% 9%. **B)** Using the *k* = 3,000 mask, MVPA classifiers trained on residual activity patterns from the background FC processing pipeline failed to differentiate task conditions.

**Supplementary Figure 3.**
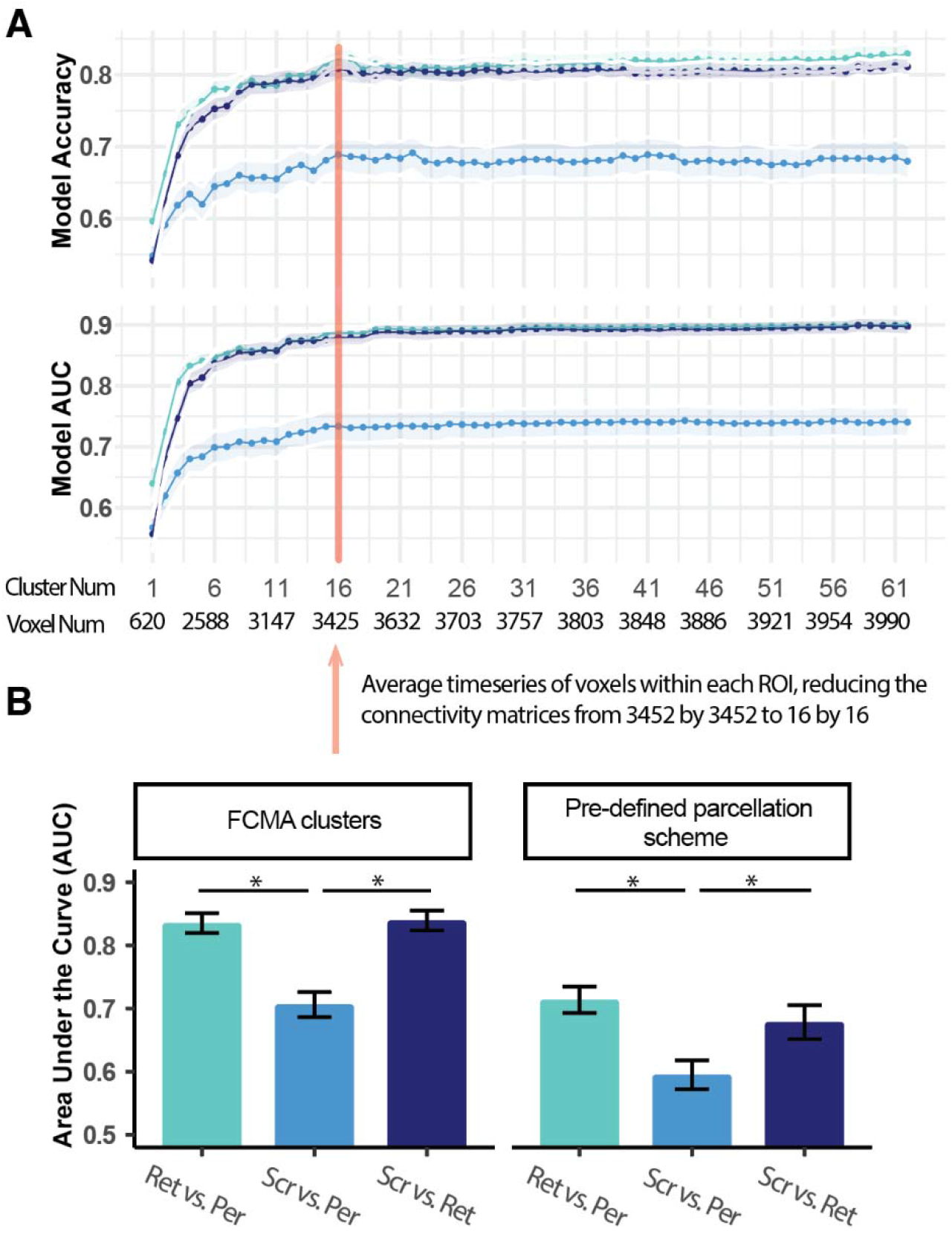
Cluster selection process. **A)** Model performance as measured by both proportion correct accuracy and area under the curve (AUC) when sequentially adding in all voxels from the next largest cluster. The semi-transparent bands indicate the standard error of the mean across all testing folds. For example, we started by using the background FC matrices of all voxels only in the largest cluster (i.e., shaped 620 x 620) to differentiate each task condition comparison. Then we added in all voxels (519; **Table S1**) from the next largest cluster and estimated the model performances of the combined background FC matrices (now shaped 1139 x 1139), and so forth. We selected a cluster number that, for either AUC or ACC, was significantly greater than the preceding number and not significantly less than the maximal number of clusters. **B**) **Left**: We averaged all voxels within a cluster, thus reducing the dimensions of the FC matrices from 3452 x 3452 to 16 x 16 and examined the performances of the models for the reduced matrices. **Right:** Instead of defining clusters using the FCMA-then-clustering pipeline, we selected the top-performing 16 parcels (from the Schaefer 1000 parcellation scheme) based on separating perception from retrieval states using the same cross-validation framework (**Figure S1**). We then examined the performance of models trained using FC matrices among predefined Schaefer parcels for differentiating each task condition comparison. The error bars represent the standard errors of the mean across all testing folds.

**Supplementary Figure 4.**
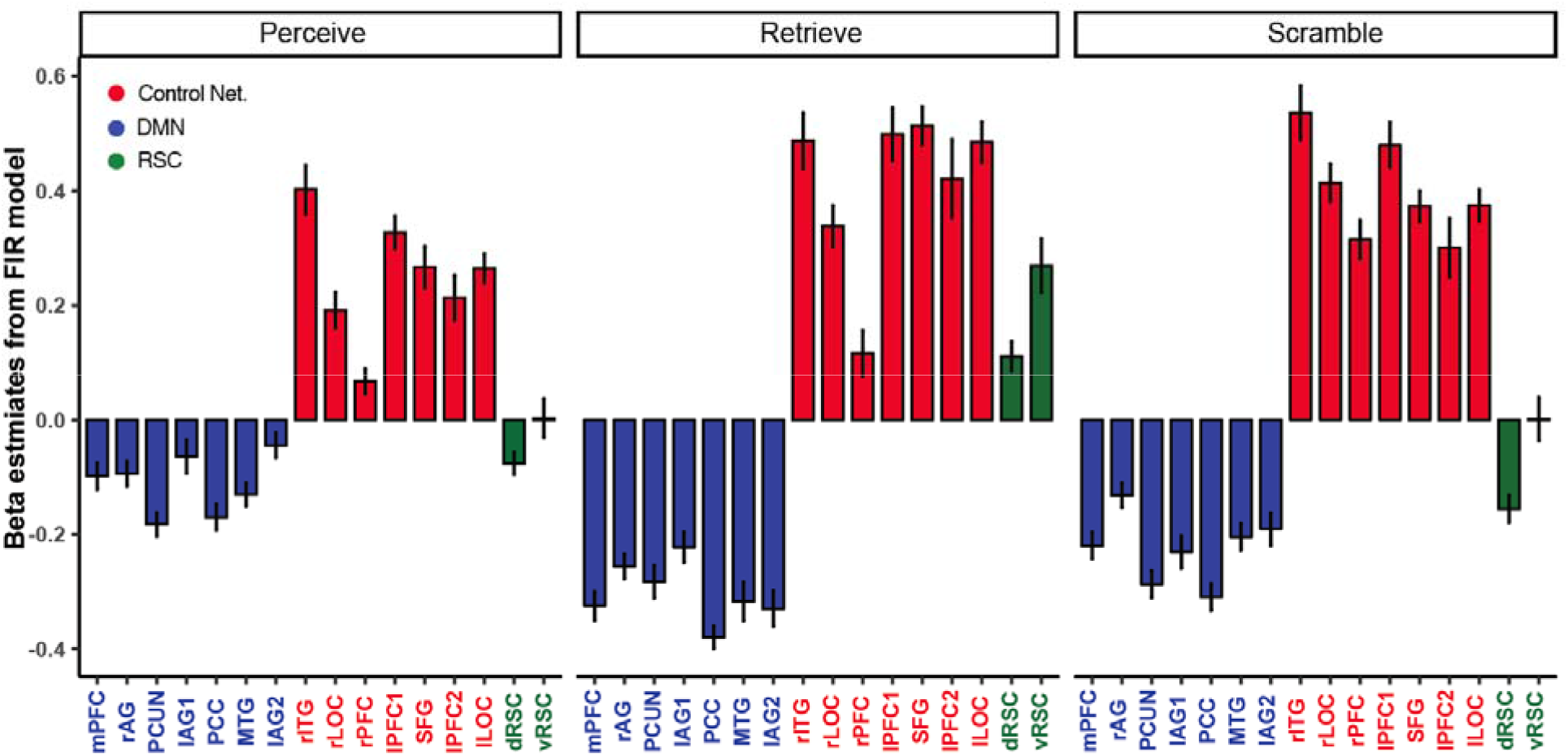
Univariate activation profiles of the 16 clusters across 3 functional communities during each task condition. The FIR model consists of 36 (4 TR instruction + 24 TR task + 8 TR inter-block interval) x 2 (epoch category) x 3 (condition) = 216 regressors. Thus, 24 (TR task) x 2 (epoch category) = 48 regressors modeled task activations for each condition. Here for each subject, we computed the averaged beta estimates (of the 48 regressors) for all voxels within a cluster per condition. Error bars indicates the standard error of the mean of beta estimates across subjects. The color indicates the functional community assignment of each cluster.

**Supplementary Figure 5.**
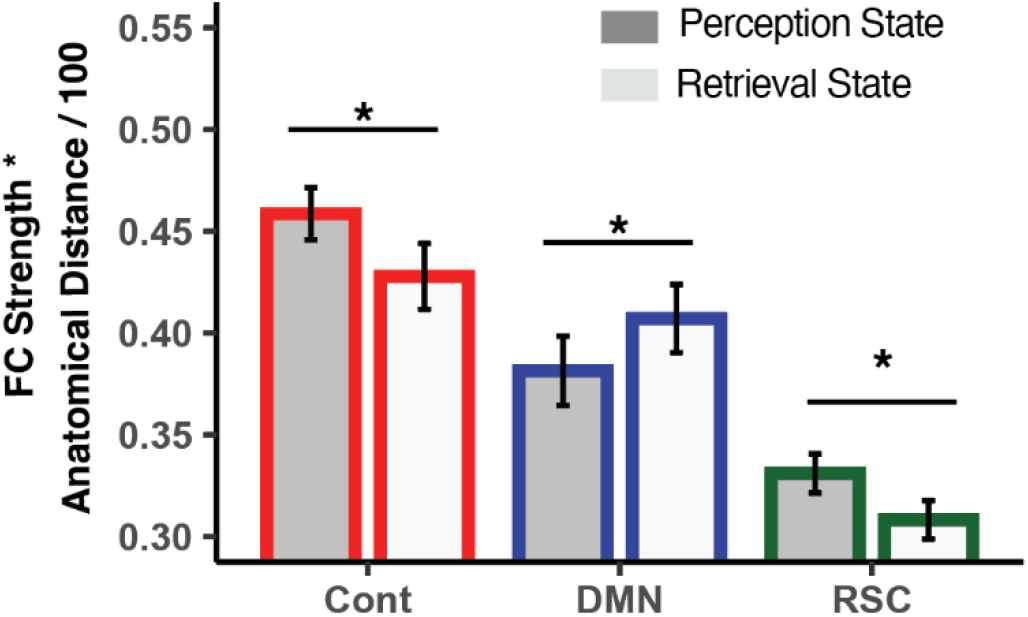
Background FC strength averaged across all pairwise connections within the same functional community across perception and retrieval states after factoring in anatomical distance. Using MNI coordinates in **Table S1**, anatomical distance was quantified as the Euclidean distance between each pair of clusters (divided by 100 to make the y-axis unit comparable to Figure 5B). The adjusted background FC strength was computed as, thus closer anatomical distance (e.g., the two RSC cluster) would lead to smaller adjusted FC strength. After factoring in anatomical distance, clusters within the Control network retained overall stronger connectivity density compared to the DMN (*t*_(23)_ = 4.03, *p* < 0.001, 95% CI = [0.02, 0.07], Cohen’s d = 0.70). On the other hand, the strong coupling strength between RSC regions observed in Figure 5B could be largely attributed to the short anatomical distance between the two RSC clusters.

**Supplementary Table 1.** Sizes and locations of the clusters of interest. The Volume column indicates the number of voxels included in each cluster. The MaxInt column indicates the maximum z-score within each cluster.

| ClusterIdx | Brain Region | Volume | X | Y | Z | MaxInt |
| --- | --- | --- | --- | --- | --- | --- |
| 1 | Medial prefrontal cortex | 620 | -0.5 | 44.3 | -1.0 | 9.14 |
| 2 | R-Inferior parietal lobule | 519 | 60.1 | -26.4 | 49.0 | 8.03 |
| 3 | Dorsal retrosplenial cortex | 400 | -18.2 | -69.3 | 24.0 | 7.88 |
| 4 | L-Intraparietal sulcus | 377 | -33.3 | -59.2 | 41.5 | 8.30 |
| 5 | R-Intraparietal sulcus | 364 | 29.8 | -66.8 | 36.5 | 7.09 |
| 6 | Precuneus | 308 | -10.6 | -54.2 | 44.0 | 7.47 |
| 7 | L-Inferior parietal lobule1 | 117 | -66.2 | -51.7 | 9.0 | 5.96 |
| 8 | R-Lateral prefrontal gyrus | 116 | 52.5 | 9.0 | 19.0 | 6.43 |
| 9 | L-Lateral prefrontal gyrus1 | 116 | -53.6 | 16.5 | 31.5 | 7.30 |
| 10 | Posterior cingulate cortex | 107 | 2.0 | -18.8 | 44.0 | 7.37 |
| 11 | Ventral retrosplenial cortex | 103 | 14.6 | -54.2 | 14.0 | 7.06 |
| 12 | Superior frontal gyrus | 82 | -3.0 | 16.5 | 51.5 | 7.27 |
| 13 | L-Lateral prefrontal gyrus2 | 64 | -48.5 | 29.2 | 24.0 | 6.23 |
| 14 | Posterior middle temporal gyrus | 58 | 62.6 | -16.3 | -11.0 | 6.30 |
| 15 | L-Inferior parietal lobule2 | 52 | -61.1 | -49.1 | 44.0 | 7.00 |
| 16 | R-Inferior temporal gyrus | 49 | 55.1 | -56.7 | -13.5 | 5.83 |

## Footnotes

1 Due to the preprint policy of biorxiv, we are unable to show the actual face stimuli used in the experiment. For added details on the stimuli, see **Methods: *Material***.

2 Note that some of the clusters (right intraparietal sulcus and the inferior temporal gyrus) had peak coordinates in the dorsal-attention network (according to Schaefer et al., 2018). Given that the remainder of the clusters (5 out of 7 clusters) fell into the Control Network we use that as the shorthand label.

3 Here we mean “visual category” to refer to the underlying stimuli used regardless of how scrambled they were. Although images were slightly scrambled, they still afforded well-off-chance M/F N/M judgements, making them dissimilar to traditional completely “scrambled” stimuli. Notably, calculating visual-category sensitivity using only non-scrambled stimuli does not change the results.

4 Note that there was overlap between the feature selection process and the final testing process (both used all subjects) in the 16 cluster analysis. Thus, the performance of the 16 clusters is slightly biased towards higher performance.

## Notes

### Competing Interest Statement

The authors have declared no competing interest.

### Summary of Updates

Changed the format to make method first. Fix a couple statistical tests to make sure all tests were performed within subjects.

